# Preserving tissue structure through density-based spatial analysis with scider

**DOI:** 10.1101/2025.09.11.675745

**Authors:** Mengbo Li, Ning Liu, Quoc Hoang Nguyen, Yunshun Chen

**Author notes:** These authors contributed equally (Mengbo Li, Ning Liu).

## Abstract

In spatial transcriptomics, most existing approaches for spatial domain detection analyze slide-wide gene expression patterns while integrating information from local neighborhoods or nearest neighbors. However, such global strategies can overlook tissue organization and obscure regional heterogeneity. To address this, we present scider, a framework for spatial data analysis to preserve tissue structures. scider defines spatial domains from local cellular organization and distribution, instead of relying on global clustering of gene expression and neighborhood profiles. Kernel density estimation (KDE) is used to characterize the spatial distribution of cells, enabling the robust and unbiased identification of regions of interest (ROIs) beyond histological annotations. We show that ROIs make multi-sample analysis possible and reveal spatial patterns that are undetectable in single-sample analysis. Additionally, KDE-derived contour lines define regions of similar cell density, supporting cell type composition and differential expression analyses along continuous spatial gradients.

## Introduction

The spatial distribution of cells in tissues is tightly controlled and closely linked to their roles in health and disease [1]. To investigate these patterns, spatially resolved omics technologies enable the molecular characterization of cellular profiles within their native tissue contexts [2]. In particular, spatial transcriptomics measures gene expression while preserving the positional context of cells in a tissue section [3]. Since its introduction, spatial transcriptomics has become a widely used tool in biomedical research, providing insights into how cells function and interact within a tissue across a range of biological contexts such as cancer and brain diseases [4, 5, 6, 7, 8].

In most spatial transcriptomics applications, the aim is to identify spatially organized cell patterns and tissue-specific gene expression signatures that inform biological processes. To this end, a fundamental step is to identify spatial domains for downstream analysis. Spatial domains refer to spatially contiguous regions within a tissue where cells share similar molecular profiles and/or are organized in a way that reflects local tissue architecture and functions. Once defined, spatial domains can serve as a basis for further analyses such as characterizing cell type composition, identifying spatially variable genes, and investigating cell-cell interactions within and between domains [9, 10, 11].

Many computational methods have been developed to identify spatial domains for spatial transcriptomics data, including SpaGCN [10], Squidpy [12], Giotto [13], BANKSY [14], and BayesSpace [15]. These approaches generally integrate gene expression within local neighborhoods to perform clustering. Each resulting cluster represents a putative spatial domain, which is then visualized post-hoc on tissue coordinates to see if it corresponds to known histological structures.

However, such an approach overlooks the underlying tissue structure. Existing methods typically annotate spatially distinct regions that share similar local features as the same spatial domain or niche, masking higher-order spatial organization patterns. For instance, in cancer, even within the same section, tumors from different regions may exhibit distinct biological behaviors and clinical outcomes due to their microenvironmental contexts. Yet, clustering often assigns multiple dispersed tumor patches to the same domain if they share similar local gene expression features, as it ignores the broader tissue architecture that distinguishes one region from another. Additionally, because clustering relies on dimensionality reduction and the selection of highly variable genes (HVGs), subtle but biologically relevant differences can easily be masked by the dominant sources of variation.

Alternatively, spatial domains can be obtained manually as regions of interest (ROIs) annotated by histologists or pathologists, such as in [11]. While manual annotations effectively preserve the tissue structure and anatomical features, they are labor-intensive, not scalable to large datasets, greatly subjective and often vary between analysts.

To this end, we propose a completely new approach for spatial domain identification determined by the spatial organization of cells. This is achieved by identifying spatial regions primarily based on the spatial coordinates of cells, as implemented in the Bioconductor R package scider. More explicitly, the default scider pipeline takes cell type annotations as input, and defines spatial regions according to the density of a cell type of interest (COI) across the slide. Kernel density estimation (KDE) is used to describe the spatial distribution of the COI, based on which regions of interest (ROIs) are identified using a graph analysis approach.

In contrast to the existing approaches where spatial information is typically used secondarily for aggregating gene expression from neighboring spots, scider uses the spatial coordinates of cells as the primary information to define spatial domains. ROIs can then be used for downstream analysis such as clustering and DE analysis by pseudo-bulking cells located in each ROI. We demonstrate in this article that ROI-based analysis helps dissect inter-tumor heterogeneity by identifying tumorspecific ROIs and comparing their gene expression profiles.

Compared to spatial domains from existing methods, ROIs allow the characterization of spatial regions at varying scales using user-defined parameters. ROIs also provide a novel framework for multi-sample analysis, enabling the joint examination of biological replicates and comparisons across groups or conditions. In contrast, when spatial domains are defined directly from gene expression data, multi-sample analysis can be more challenging due to batch effects and the need for normalization. Compared to manual annotations, scider performs automatic ROI detection that is both reproducible and scalable. Compared to clustering-based algorithms, scider identifies COI-specific ROIs within seconds, offering substantially greater computational efficiency while crucially preserving the structural organization of COI-specific regions across the tissue.

In complement to ROI-based analyses, spatial regions can also be defined as contour regions based on the spatial density of the COI. Contour-based analysis enables the investigation of cellular and molecular patterns along the spatial gradient of COI density within specific microenvironments. For example, one application of contour-based analysis is to examine immune cell infiltration by assessing cell type composition across tumor density levels.

## Results

### scider defines a novel framework for spatial transcriptomics data analysis

scider provides two complementary approaches to identify spatial regions for analysis of spatial transcriptomics data, that is, regions of interest (ROIs) and contour regions (Figure 1A). With designated spatial regions, scider offers a streamlined suite of tools for data exploration and visualization. For downstream analysis, scider integrates seamlessly with limma and edgeR pipelines by generating pseudo-bulk samples from cells within each ROI and/or contour region. This allows flexible experimental designs at both ROI- and contour-levels to address a broad range of biological questions.

**Figure 1.**
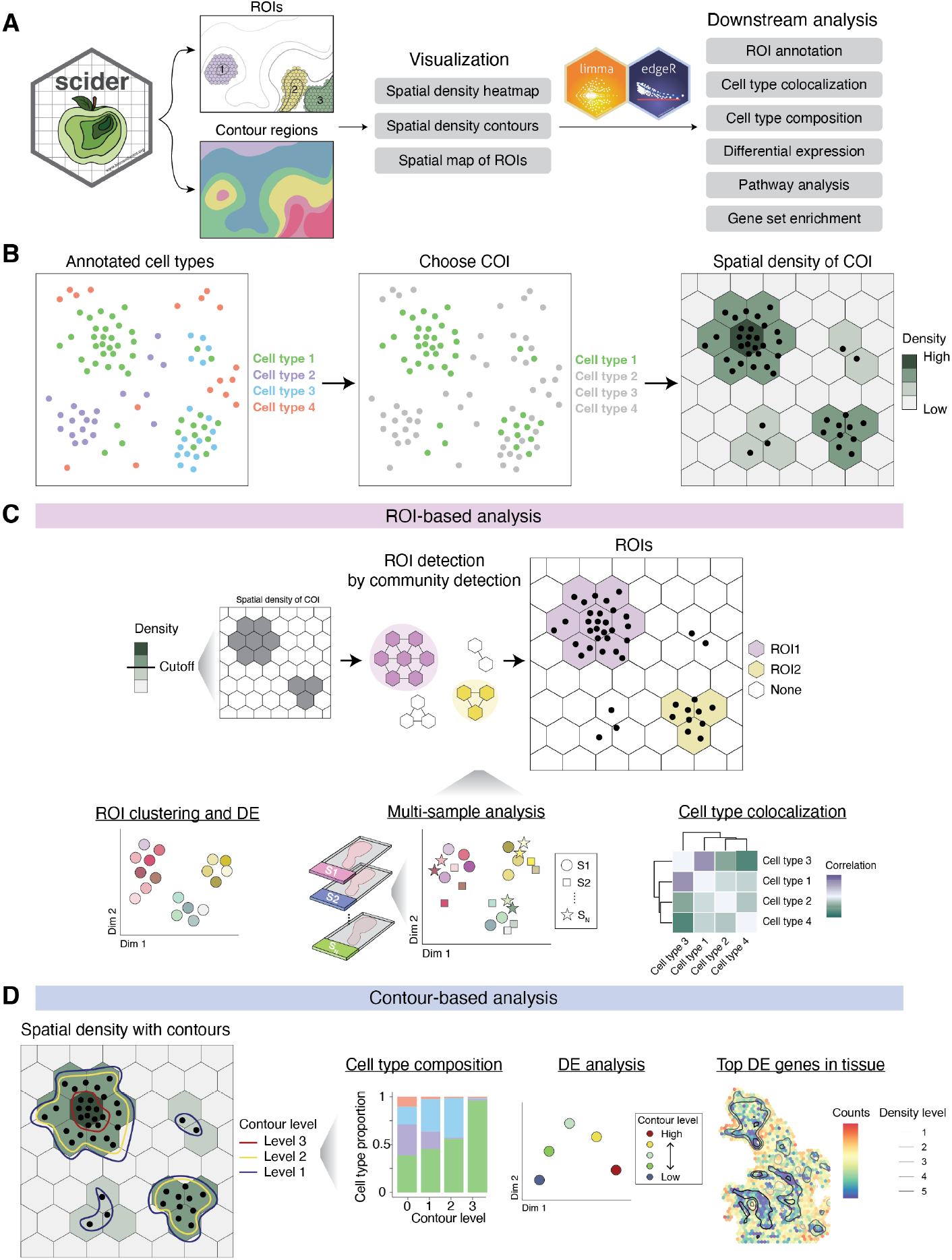
Overview of scider workflow. (A) Summary of scider functionalities. (B) A typical scider analysis takes spatial transcriptomics data with cell type annotation as input. A cell type of interest (COI) is chosen. Spatial density of the COI is estimated by dividing the slide into grids (rectangles or hexagons). (C) Region of interest (ROI)-based analysis. ROIs are identified by creating a graph based on the spatial density of the COI as described in Methods. Given the ROIs, downstream analysis such as clustering, differential expression (DE) and multi-sample analysis can be performed by pseudo-bulking cells in each ROI. Cell type colocalization analysis can also be performed on the ROI level. (D) Contour-based analysis. Contours are calculated at a number of density levels based on the spatial density function. Cells are assigned to each density level of the COI for downstream analysis and visualization.

Briefly, a typical scider workflow defines spatial regions based on the spatial density of a cell type of interest (COI) (Figure 1B). Given user-provided cell type annotations, a COI can be chosen for downstream analysis. scider computes the spatial density of the COI using kernel density estimation (KDE). The estimation is performed by mapping COI cells onto rectangular or hexagonal grids spanning the whole slide.

The first approach is to identify COI-specific regions of interest (ROIs) (Figure 1C). Given the spatial density of the COI, scider identifies ROIs by constructing a weighted and undirected graph after filtering out grids of COI densities below a user-defined threshold. To construct the graph, nodes are represented by grids, and edges are defined by the physical adjacency between grids. Edges are weighted by the average COI density between pairs of adjacent grids. The graph is then partitioned into communities using community detection algorithms, where each community corresponds to an ROI. Depending on the nature of the sample, ROIs can be discrete or continuous, and can vary in size. To accommodate heterogeneous samples, ROI detection is fully automated and can be tuned by adjusting the grid size, bandwidth and community detection method.

Given the ROIs, scider unlocks a range of ROI-based analyses such as clustering, differential expression (DE) and multi-sample analysis by pseudo-bulking cells in each ROI. Cell type colocalization analysis can be performed on both ROI and whole-slide levels, where colocalization is quantified by the Pearson’s correlation coefficient between the spatial densities of two cell types within the spatial region.

The second approach analyzes contour regions derived from the COI density (Figure 1D). Contour lines are calculated at a number of density levels based on the spatial density function, after which cells are assigned to each density level of the COI. Cell type composition can then be visualized at each contour level. For instance, this approach can be applied to characterize immune cell infiltration at varying tumor density levels . DE analysis can also be performed by pseudo-bulking cells at each contour level. Contour-based analysis enables the investigation of cellular and molecular patterns along the spatial gradient of COI density within specific microenvironments. This allows us to zoom in on specific microenvironments for deeper insights into the localized cellular organization and functions.

### scider identifies cell type-specific ROIs

A standard scider workflow is demonstrated in the next few sections using a public Xenium *in situ* breast carcinoma data. Two serial 5 *μ*m FFPE tissue sections were obtained from the same breast cancer block (replicate 1 and replicate 2 of Sample #1, ER^+^/HER2^+^/PR^-^) [11]. We obtained cell type annotations from the original study [11]. Several cell types were merged for simplicity (as summarized in Methods). Figure 2A shows the spatial plot with updated cell type annotations, and Supplementary Figure 1 shows the spatial plot of each individual cell type. For ROI-based analyses, the cell-type of interest (COI) was chosen to be the combination of ductal carcinoma *in situ* (DCIS) and invasive tumor cells.

**Figure 2.**
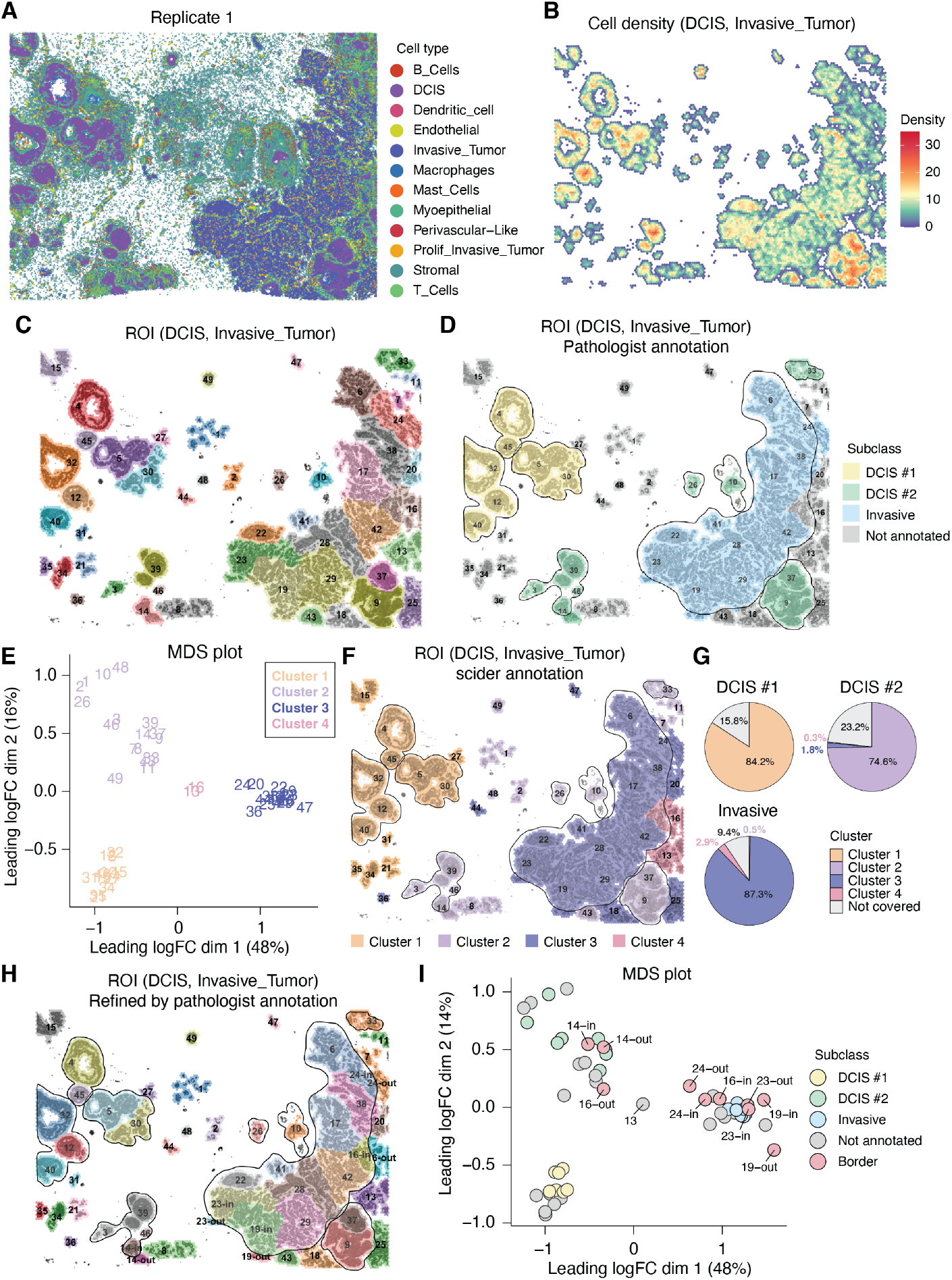
ROI detection on a Xenium breast carcinoma sample (replicate 1) compared with its pathologist annotation. DCIS and invasive tumor cells are used as the COI. (A) Spatial plot with cell type annotation. (B) Heatmap of spatial density of DCIS and invasive tumor cells. Median of the estimated densities of all grids is calculated. Grids of densities less than the median are filtered out in this visualization. (C) ROIs detected by scider. DCIS and invasive tumor cells are highlighted in the background, similarly for panels D, F and H. (D) scider ROIs overlaid by pathologist-annotated ROIs from [11]. (E) MDS plot of gene expression in COI cells on the ROI level by scider. (F) scider ROIs colored by clusters based on the MDS plot (panel E), overlaid with pathologist annotations. (G) Percentage area overlapped by scider ROIs for each pathologist-annotated ROI. (H) scider ROIs refined by pathologist annotations. scider ROIs at the border of pathologist-annotated ROIs are divided into two subareas, one inside the pathologist-annotated region and the other outside. (I) MDS plot of gene expression in COI cells across scider ROIs refined by pathologist annotation from panel H.

The spatial density of the COI was estimated by dividing the slide into 50 *μ*m-wide hexagonal grids for kernel density estimation (KDE). Figure 2B visualizes the estimated spatial density of the COI, where grids with low COI densities (less than 0.5) were filtered out. A weighted and undirected graph was constructed based on the spatial density of the COI, and regions of interest (ROIs) were subsequently identified via community detection. Grids of each community comprise an ROI as illustrated in Figure 2C. To focus on regions of sufficient size, ROIs with less than 20 grids were filtered out.

scider identifies cell type-specific ROIs based on the spatial density of the COI. ROI calculations are fully automated and can be tuned to accommodate heterogeneous samples by adjusting the grid size, bandwidth and community detection method. Supplementary Figure 3 shows the effect of varying bandwidth values on ROI detection results. At a fixed grid size, smaller bandwidths typically produce ROIs with finer spatial resolution. ROI identification is completely reproducible given the same parameters.

For benchmarking, we compared scider-detected ROIs with the histology/pathology and scFFPEseq guided ROIs for DCIS#1, DCIS#2 and invasive tumor from the same study [11]. For simplicity, we refer to these ROIs as ‘pathologist-annotated ROIs’ hereafter. Figure 2D depicts the pathologist annotations overlaid on top of the scider ROIs. Each scider ROI is annotated as the corresponding pathologist ROI if the percentage of its area overlapped by the pathologist ROI is greater than 50%. Overall we observed substantial agreement between the two sets of ROIs. A few scider ROIs lack corresponding pathologist annotations, because the manual annotations did not encompass all areas containing the COI cells.

We next derived annotations for scider ROIs based on gene expression data. Pseudo-bulk samples were created by aggregating COI cells within each ROI and visualized by the multi-dimensional scaling (MDS) plot (Figure 2E). Four clusters of ROIs were identified by visual inspection of the MDS plot. When mapped back onto the spatial plot, the four ROI clusters showed great concordance with different pathologist-annotated ROIs of the sample. In particular, cluster 1 primarily aligns with the DCIS#1 region, cluster 2 with the DCIS#2 region and cluster 3 with the invasive tumor region (Figure 2F). Notably, we saw greater variation in the MDS plot between ROIs of cluster 2 compared to those of clusters 1 and 3. As seen in the spatial plot, ROIs from clusters 1 and 3 were more spatially contiguous, whereas ROIs from cluster 2 were more dispersed across the slide. Meanwhile, in the MDS plot, ROIs 13 and 16 were located near the intersection of clusters 1, 2 and 3. These two ROIs lied at the interface of DCIS#2 and invasive tumor regions, and without further analysis, we kept them as their own cluster for now. This analysis shows that scider ROIs can be easily annotated using gene expression data, independent of pathologist input. This enables the inclusion of additional regions beyond those manually annotated by pathologists for downstream analyses.

We then quantified the overlap between pathologist-annotated and scider ROIs by computing the proportion of each pathologist-annotated ROI that intersected with any scider ROI (Figure 2G). All pathologist-annotated ROIs showed substantial overlap (over 80%) with scider ROIs, except for the DCIS#2 region. The DCIS#2 region had lower coverage by scider ROIs because we had filtered out the smaller ROIs near ROI 10. We could fine-tune our results to increase the overlap with pathologist-annotated ROIs by retaining the smaller ROIs or using larger bandwidths. However, we chose to keep the results as is to demonstrate a typical scider workflow. Moreover, recovering pathologist-annotated ROIs is neither the ultimate goal nor a definitive benchmark for ROI detection, as these annotations are often subjective.

Finally, we examined the scider ROIs that were split at the border of pathologist-annotated ROIs. Based on gene expression data, we aimed to assess if the boundaries drawn by pathologists were definitively grounded in biological relevance, or if they exhibited some degree of arbitrariness. We split a few scider ROIs at the border of pathologist-annotated ROIs into two subareas, one inside the pathologist-annotated region and the other outside (Figure 2H). Pseudo-bulk samples were created by aggregating COI cells within each refined ROI and visualized by the MDS plot as Figure 2I. Most of the subareas of the same ROI were clustered together, except for ROI 16. This confirms that while pathologist-annotated ROIs are generally biologically meaningful, the delineation of their boundaries is often influenced by subjective judgement. In contrast, scider provides a more objective, datadriven, and reproducible approach to identify ROIs.

### ROIs provide a pathway to multi-sample analysis

Importantly, ROIs define a common framework for jointly analyzing biological replicates and comparing samples across different groups or conditions. Here, we applied scider to the two Xenium *in situ* replicates of Sample 1, where the COI was still the combination of DCIS and invasive tumor cells. The spatial plots of the two replicates are shown in Supplementary Figure 3A and B, with the same cell type annotations as in Figure 2A. The spatial density of the COI was estimated for each replicate and visualized by the heatmaps in Supplementary Figure 3C and D. ROIs were identified as described in the previous section and illustrated in Supplementary Figure 4A. We saw that scider was able to identify consistent ROI patterns between the two serial section replicates. For example, ROIs in the DCIS#1 region were repeatedly identified across both replicates with similar spatial locations and shapes, even when partially cropped in replicate 2 (e.g., ROIs 34 and 36).

We proceeded to annotate the scider ROIs using gene expression data, as described in the previous section. For each replicate, pseudo-bulk samples were created by aggregating COI cells within each ROI and visualized by the MDS plot in Figure 3A and Supplementary Figure 4B. We observed no clear batch effects between the two replicates. Four clusters of ROIs were identified by k-means clustering, which were then mapped back onto the spatial plots (Figure 3B). The two replicates demonstrated strong concordance, with ROIs in each cluster consistently occupying the same regions between the two serial sections. ROIs from cluster 1 primarily align with the DCIS#1 region, cluster 2 with the DCIS#2 region and cluster 3 with the invasive tumor region, similar to the results in Figure 2F. Cell type composition analysis also confirmed our annotation of the ROI clusters (Figure 3C, left). Previously, we were not able to confidently assign ROIs 13 and 16 of replicate 1 to any cluster when replicate 1 was analysed alone. However, joint analysis with replicate 2 allowed us to assign them to DCIS#2 (cluster 2).

**Figure 3.**
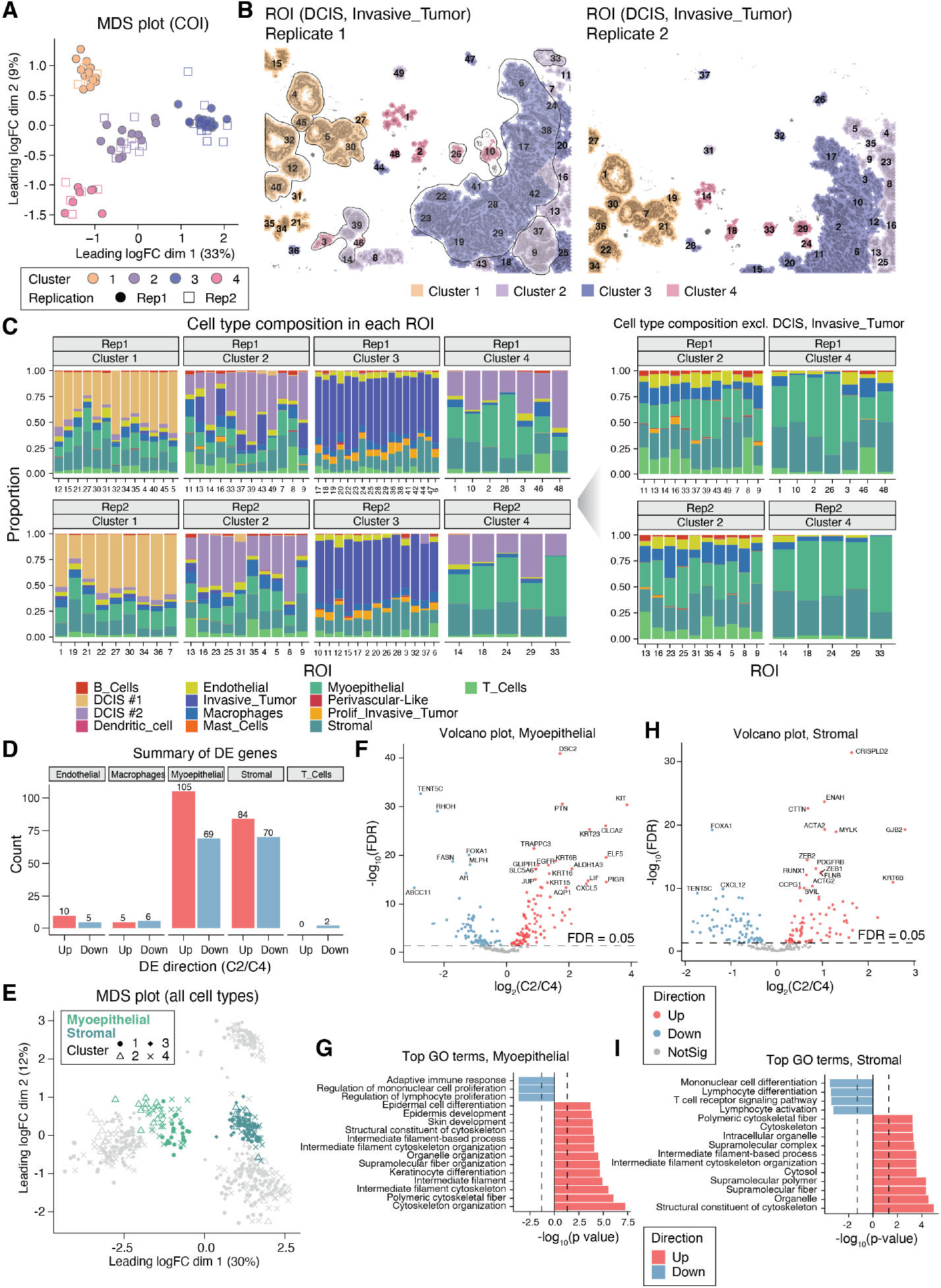
Joint analysis of replicates of the breast carcinoma sample by Xenium leveraging the ROIs. DCIS and invasive tumor cells are used as the COI. (A) MDS plot of gene expression in COI cells from ROIs in both replicates. (B) Spatial plots of scider ROIs colored by clusters identified in the MDS plot (panel A) for replicate 1 (left) and replicate 2 (right). (C) Cell type composition of each ROI with the COI included (left) or excluded (right). (D) Summary of DE genes for each cell type comparing those in ROIs of cluster 2 (C2) with those in cluster 4 (C4). (E) MDS plot of gene expression in all cell types from ROIs in both replicates. Myoepithelial and stromal pseudo-bulk samples are highlighted. Shapes correspond to the ROI clusters identified in panel A. The same MDS plot with all cell types annotated is available as Supplementary Figure 4B. (F) Volcano plot comparing myoepithelial cells from C2 ROIs with those of C4 ROIs. (G) Top GO terms comparing myoepithelial cells from C2 ROIs with those of C4 ROIs. (H) Volcano plot comparing stromal cells from C2 ROIs with those of C4 ROIs. (I) Top GO terms comparing stromal cells from C2 ROIs with those of C4 ROIs.

Interestingly, we also observed a fourth cluster of ROIs (cluster 4) that were not present in the previous analysis. Both clusters 2 and 4 had substantial proportions of DCIS#2 cells. In contrast to cluster 4, several cluster 2 ROIs (e.g., ROIs 13 and 16) contained a prominent proportion of invasive tumor cells (Figure 3C, left). Upon excluding the COI cells, cluster 4 ROIs were found to be primarily composed of myoepithelial and stromal cells, whereas cluster 2 ROIs exhibited a more heterogeneous cellular composition (Figure 3C, right). Compared to cluster 2, cluster 4 ROIs had higher proportions of several other cell types, most notably endothelial cells, macrophages and T cells.

To further investigate the differences between clusters 2 and 4, we performed differential expression (DE) analysis to compare each cell type between the two clusters. We first compared the tumor cells in clusters 2 and 4, where pseudo-bulk samples were created by aggregating COI cells within each ROI. We identified 68 upregulated and 78 downregulated genes in cluster 2 compared to cluster 4 at 5% FDR (Supplementary Figure 4C, right). Cluster 2 regions had higher expression of structural and epithelial markers including KRT5, KRT8, KRT23 and PTN [16]. Cluster 4 showed the upregulation RHOH and TENT5C, which is related to immune signaling [17] and RNA regulation [18], respectively (Supplementary Figure 4E). Consistently, GO analysis (Supplementary Figure 4G) showed that cluster 4 regions were enriched for immune-related processes such as leukocyte and lymphocyte activation, whereas cluster 2 regions were associated with cytoskeletal organization and intermediate filament-related pathways. Notably, regions near ROIs 3, 10, 26, and 46 were annotated as DCIS#2 regions by pathologists [11], yet we showed that molecularly, they are distinct from the other DCIS#2-enrich regions of cluster 2.

Next, pseudo-bulk samples were created by aggregating cells within each ROI for every cell type and visualized by the MDS plot in Supplementary Figure 4H. We observed distinct clustering by cell type with no evident batch effect between the two replicates. The same MDS plot was annotated by ROI cluster in Figure 3E, with the myoepithelial and stromal pseudo-bulk samples highlighted. In particular, there was clear separation of myoepithelial cells across the four ROI clusters.

Figure 3D summarizes the DE results for each cell type, where myoepithelial cells and stromal cells had the most significant DE genes between ROIs of clusters 2 and 4. In myoepithelial cells, the DE analysis identified 105 upregulated genes and 69 downregulated genes in cluster 2 compared to cluster 4 at 5% FDR (Figure 3F). KIT, ELF5, CLCA2, and CXCL5 were among the top upregulated genes in cluster 2, whereas FOXA1, AR and FASN were more highly expressed in cluster 4. This expression pattern suggested that ROIs from cluster 2 were enriched for basal-like, progenitor-associated signatures, while cluster 4 ROIs were more consistent with a luminal-like, differentiated phenotype. Such an observation is also consistent with results from the cell type composition analysis, where we observed reduced infiltration of endothelial cells, macrophages, and T cells in cluster 4 compared to cluster 2 [19, 20]. We further investigated the DE genes by performing a gene ontology (GO) analysis (Figure 3G). GO terms such as cytoskeleton organization, keratinocyte differentiation and epidermis and skin development were upregulated in cluster 2, corresponding to basal epithelial cell functions. Despite the upregulation of innate immune genes such as CXCL5, the top downregulated GO term suggested suppressed adaptive immune engagement in myoepithelial cells of cluster 2 compared to cluster 4.

In stromal cells, we identified 84 upregulated genes and 70 downregulated genes in cluster 2 compared to cluster 4 at 5% FDR (Figure 3H). Upregulated genes included markers of myofibroblast activation (e.g., ACTA2, ZEB1/2), cytoskeletal remodeling, and cell motility (ENAH, CTTN, FLNB), indicating a pro-invasive stromal phenotype in cluster 2 ROIs. Meanwhile, the significant downregulation of CXCL12 and TENT5C suggested a loss of stromal homeostatic and immune-regulatory functions, further supporting a tumor-promoting microenvironment. Consistent with these findings, GO analysis (Figure 3I) showed reduced lymphocyte differentiation and T cell signaling, along with increased cytoskeletal organization and structural cytoskeleton components, indicating a shift toward a contractile, immune-suppressive stromal state in cluster 2.

Together, these results suggest that cluster 2 regions may represent a more progression-prone subtype. These findings also highlight the molecular heterogeneity within histologically-annotated DCIS#2 regions.

### Colocalization analysis identifies cell type patterns across spatial regions

ROIs also enable cell type colocalization analysis. Cell type colocalization measures how frequently two cell types occur together within the same spatial region. In scider, it is quantified using a Pearson’s correlation coefficient between the spatial densities of the two cell types within a spatial region. Positive correlations indicate frequent co-occurrence, while negative correlations suggest mutual spatial exclusion.

In the following sections, we focus on invasive tumor cells as the COI in the Xenium *in situ* breast carcinoma sample (replicate 1). The spatial density of invasive tumor cells was estimated on a 50 *μ*m-wide hexagonal grid and visualized by the heatmap in Supplementary Figure 6. Grids with an estimated density less than 0.5 were filtered out. ROIs were detected as described in a previous section, and ROIs with less than 50 grids were filtered out. Figure 4A shows the spatial plot of invasive tumor ROIs with invasive tumor cells overlaid in the background. We then performed cell type colocalization analysis on the ROI level.

**Figure 4.**
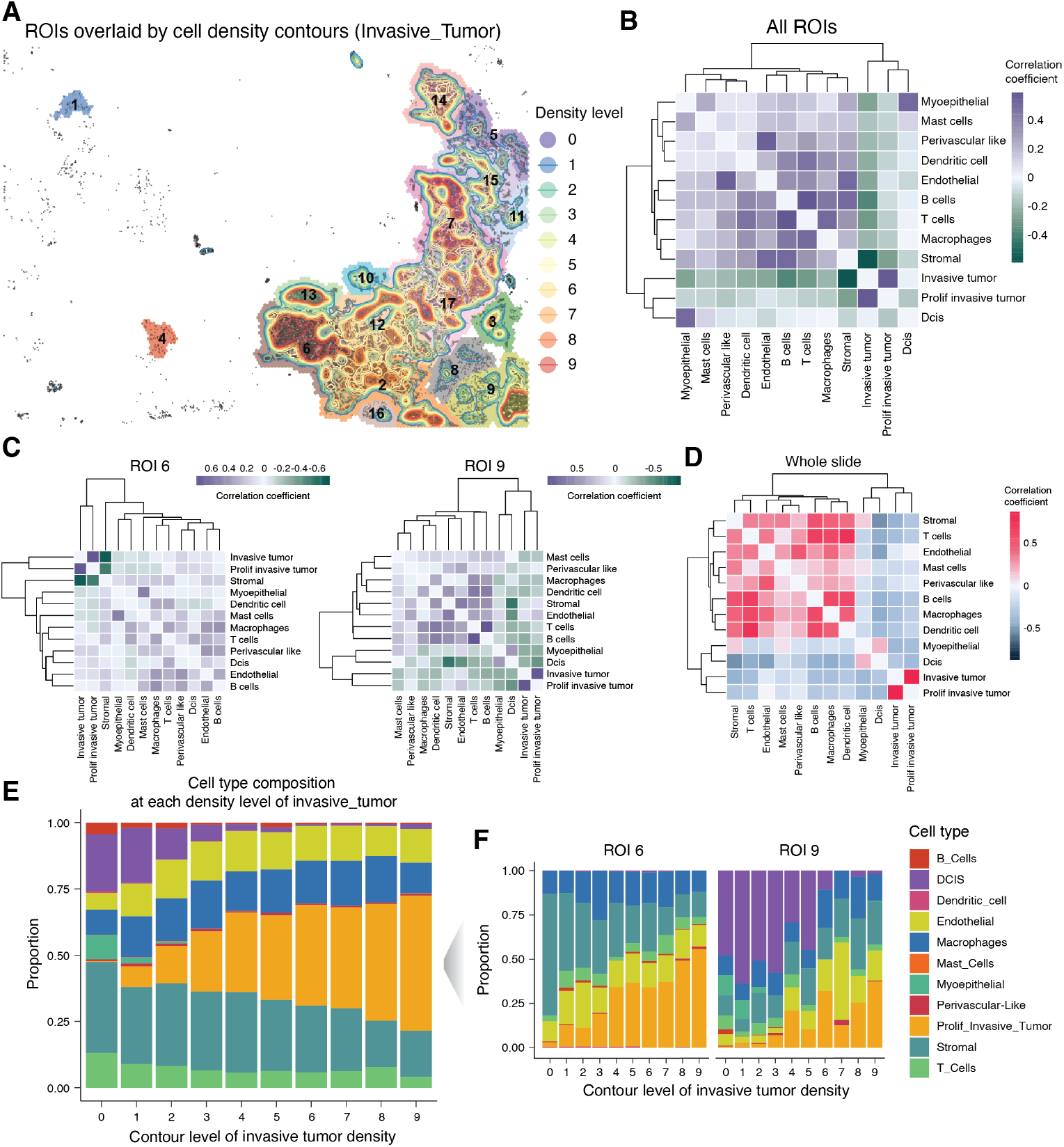
Cell type colocalization and cell type composition analysis on the Xenium breast carcinoma sample (replicate 1), COI being invasive tumor. (A) Spatial plot of invasive tumor ROIs, overlaid by contour lines calculated from the spatial density of invasive tumor cells. Invasive tumor cells are shown in the background. (B) Cell type colocalization across all invasive tumor ROIs. (C) Cell type colocalization in ROI 6 (left) and ROI 9 (right). (D) Global cell type colocalization across the whole slide. (E) Cell type composition at each contour level of invasive tumor density across all ROIs. (F) Cell type composition at each contour level of invasive tumor density in ROI 6 (left) and ROI 9 (right).

Figure 4B shows the colocalization matrix across all invasive tumor ROIs. We observed distinct clustering of tumor-associated cell types (invasive tumor, proliferative invasive tumor and DCIS) separate from all other cell types. In particular, DCIS strongly co-occurred with myoepithelial cells. Invasive tumor cells showed strong colocalization with proliferative invasive tumor, but were negatively colocalized with stromal cells. Supplementary Figure 7 presents the correlation coefficients for all cell type pairs across individual ROIs, revealing broadly consistent clustering patterns. However, we saw that the colocalization patterns were ROI-specific. For example, DCIS cells were positively correlated with myoepithelial cells in most ROIs, but showed negative correlations in ROIs 4 and 8 (Supplementary Figure 7).

To further illustrate this region-specific variability, Figure 4C displays the colocalization matrix in two representative ROIs, ROI 6 and ROI 9. From previous analyses, we know that ROI 6 is primarily composed of invasive tumor cells (near ROI 23 in Figure 3B), while ROI 9 is at the interface of DCIS#2 and invasive tumor ROIs (near ROIs 9 and 25 in Figure 3B). In ROI 6 (Figure 4C, left), invasive tumor cells were highly colocalized with proliferative invasive tumor cells, while both showed strong spatial segregation from stromal cells. All other cell types exhibited weak colocalization with each other. Meanwhile, ROI 9 (Figure 4C, right) displayed a colocalization pattern more closely aligned with the overall average across all ROIs (Figure 4B). For instance, DCIS cells in ROI 9 weakly colocalized with myoepithelial cells but were negatively correlated with all other cell types. T cells showed strong colocalization with B cells, consistent with the average ROI pattern. Supplementary Figure 8 shows the colocalization patterns in each ROI. Again, it was evident that the colocalization patterns were highly context-dependent and greatly differed across different invasive tumor ROIs.

Lastly, Figure 4D visualizes the global cell type colocalization across the whole slide. Global colocalization patterns generally resembled the ROI-average. Stromal cells were negatively correlated with tumor-associated cell types but positively correlated with most others. Invasive tumor cells were strongly colocalized with proliferative invasive tumor cells, but not with DCIS or any other cell type. However, the clustering patterns differed slightly. DCIS and myoepithelial cells were more closely associated with each other than with invasive tumor and proliferative invasive tumor cells. Collectively, these results suggest that colocalization was strongly influenced by their spatial context, and can vary significantly across different spatial regions.

### Spatial density contours uncover regional variation in cell type composition

Based on spatial density estimation, scider also calculates contours at varying density levels. These contour lines define regions with similar COI density, providing an alternative approach to identify and construct spatial regions for downstream analysis. In contrast to ROIs, this approach helps elucidate microenvironmental profiles driven by continuous spatial variation in COI density, rather than by discrete regions.

With replicate 1, we continued to investigate invasive tumor cells as the COI. Given the spatial density estimation, contour lines were calculated at 0.1 density intervals, spanning values from 0.1 to 1.0. These contour lines were then overlaid on the spatial plot of invasive tumor ROIs (Figure 4A), where higher density levels correspond to regions with more invasive tumor cells. The contour lines divide the slide into 10 regions, each representing a distinct contour level. For example, level 9 corresponds to the highest-level contour, portraying localized areas with the greatest invasive tumor cell density on the slide. Cells were assigned to a contour region based on their spatial location, cell type composition were then examined at each contour level.

Figure 4E visualizes the cell type composition at each contour level across all ROIs. The proportion of proliferative invasive tumor cells increased with contour level, peaking at level 9. In contrast, DCIS proportions rapidly declined at lower levels and were nearly absent at higher contours. Stromal cells were slightly more abundant at lower contour levels, while endothelial cells, T cells, and macrophages remained relatively stable across all contours above 0. This pattern indicated a shift in the COI microenvironment from a more heterogeneous composition at lower-level contours to a more tumor-dominant profile at higher-level contours.

We next examined the cell type composition at each contour level in the two representative ROIs (ROI 6 and ROI 9; Figure 4F). The proportion of proliferative invasive tumor cells in ROI 6 was consistently higher than in ROI 9 across all contour levels. DCIS cells were nearly absent in ROI 6, while ROI 9 showed a substantial DCIS presence at lower contour levels, which declined sharply from level 6 onward. Stromal cells were more abundant in ROI 6 at lower contour levels, but their proportion decreased at higher levels. Macrophage proportions in ROI 6 remained stable across all contour levels. In contrast, ROI 9 exhibited a more variable composition across contour levels, with a slight increase in macrophages and stromal cells at higher contours.

Additionally, Supplementary Figure 9 displays the cell type composition across contours in each invasive tumor ROI. We saw distinct patterns of cell type composition across different ROIs. This contour-based analysis captures how cell type composition varies with COI density in different spatial regions.

### Contour-based DE analysis characterizes microenvironment-specific cell states

We next performed DE analysis across contour levels to identify genes associated with increasing invasive tumor cell density. Less contour levels were used in this analysis to ensure sufficient cell numbers for pseudo-bulking. Meanwhile, to aid interpretation, contour levels were defined based on the expected number of invasive tumor cells per grid at each level (see Methods for details). Figure 5A shows the spatial plot of invasive tumor cells overlaid by the new contour lines. Pseudo-bulk samples were created by aggregating cells at each contour level for all cell types.

**Figure 5.**
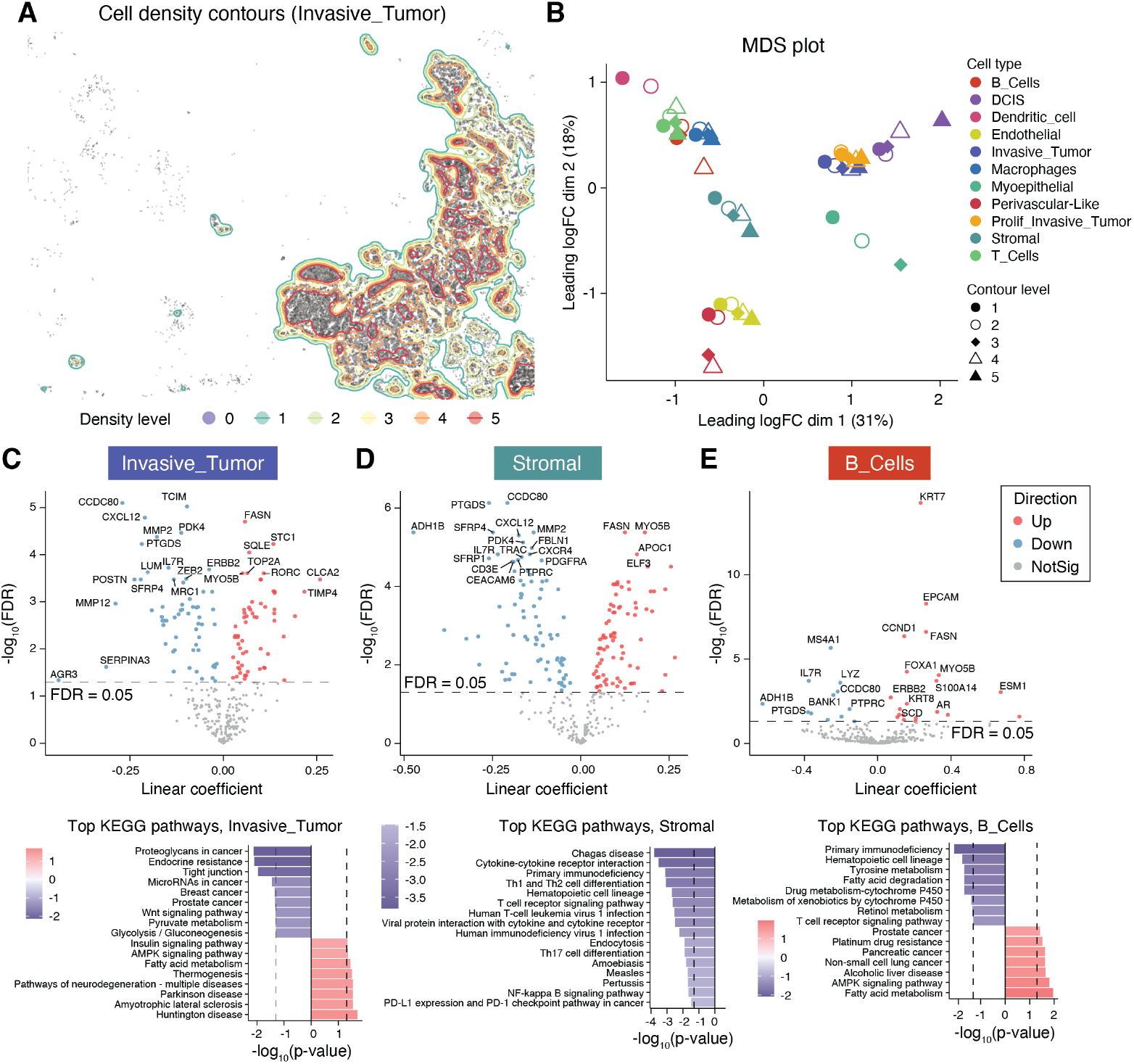
Contour-based DE analysis on the Xenium breast carcinoma sample (replicate 1), COI being invasive tumor. (A) Spatial plot of invasive tumor cells, overlaid by contour lines calculated from the spatial density of invasive tumor. (B) MDS plot of gene expression in all cell types at each contour level. Pseudo-bulk samples are created by aggregating cells at each contour level for all cell types. (C - E) Volcano plots (top) for gene expression changes and top KEGG pathways (bottom) associated with invasive tumor densities in invasive tumor cells (C), stromal cells (D) and B cells (E).

Figure 5B shows the MDS plot of gene expression across all cell types at each contour level. Clear separation of cell types was observed. Within a few cell types, such as endothelial and stromal cells, a clear trajectory emerged with increasing contour level, suggesting a transcriptomic shift in cell state associated with higher invasive tumor density.

DE analysis was performed to compare invasive tumor, stromal cells and B cells (Figure 5 C–E) located at different levels of invasive tumor density. In invasive tumor cells, genes involved in proliferation, metabolism, and extracellular matrix (ECM) remodeling, such as TOP2A, FASN, and TIMP4, were upregulated in high-density regions (Figure 5C, top). In contrast, key genes including ERBB2, CXCL12, and MMP2 were downregulated, indicating reduced HER2 signaling and altered immune or stromal interactions [21, 22, 23]. These findings suggest a shift toward a more proliferative and metabolically active invasive phenotype as tumor density increased. KEGG pathway analysis (Figure 5C, bottom) further supported this. Increasing invasive tumor density was associated with downregulation of cancer-related signaling pathways, including Wnt signaling, proteoglycans in cancer and endocrine resistance. Conversely, genes involved in fatty acid metabolism, insulin signaling and AMPK signaling were upregulated as invasive tumor density increased. This suggests a metabolic reprogramming toward lipid utilization and stress adaptation in high tumor-density regions. Furthermore, we observed downregulation of the tight junction pathway with increasing invasive tumor density, suggesting increased invasiveness at the tumor front relative to the tumor core. This is further supported by the GO analysis (Supplementary Figure 10A), which showed downregulation of terms related to locomotion, regulation of cell-substrate adhesion and cell migration.

Stromal cells showed upregulation of FASN (Figure 5D, top), indicating altered metabolic pathways supporting tumor progression [24]. In contrast, downregulation of genes such as CXCL12, MMP2, CD3E, and PTPRC suggested impaired immune signaling, ECM remodeling, and immune surveillance [25, 26, 27, 28]. These changes reflect stromal adaptation to higher tumor density, promoting a more proliferative and immune-suppressive microenvironment. The downregulation of immune-related KEGG pathways (Figure 5D, bottom), such as T cell receptor signaling and cytokine interactions, further indicated reduced immune activity. These results collectively imply a shift toward immune evasion as invasive tumor density increased.

In B cells, we observed downregulation of genes such as MS4A1, IL7R and PTPRC (Figure 5E, top). This indicated a potential impairment of immune signaling and B cell function [28, 29, 30]. The top downregulated KEGG pathway (Figure 5E, bottom) was primary immunodeficiency, which further supported the notion of reduced immune activity.

### scider models densities in sequencing-based spatial transcriptomics data

The application of scider is not limited to image-based or cancer-focused spatial transcriptomics. To demonstrate scider’s broader utility, we applied its spatial density analysis workflow to a Visium HD profile of adult mouse brain (10x Genomics, 2024, March 30). In contrast to the high-entropy topography of tumor samples, the major brain domains are visually distinct and readily resolved by cell type labels alone (Figure 6A). After selecting six spatially distinct domains, including the cortex layer 2/3 and 6, dentate gyrus, hippocampus, immune infiltrates and oligodendrocytes, scider estimated their spatial densities under equal weighting and revealed a cell density peak in the hippocampal formation (Figure 6B). Subsequently, We used this hippocampal density map to define five concentric “hippocampus-contour” regions (levels 0-4, Figure 6C), where level 0 contains virtually no hippocampal cells and level 4 is almost exclusively hippocampus.

**Figure 6.**
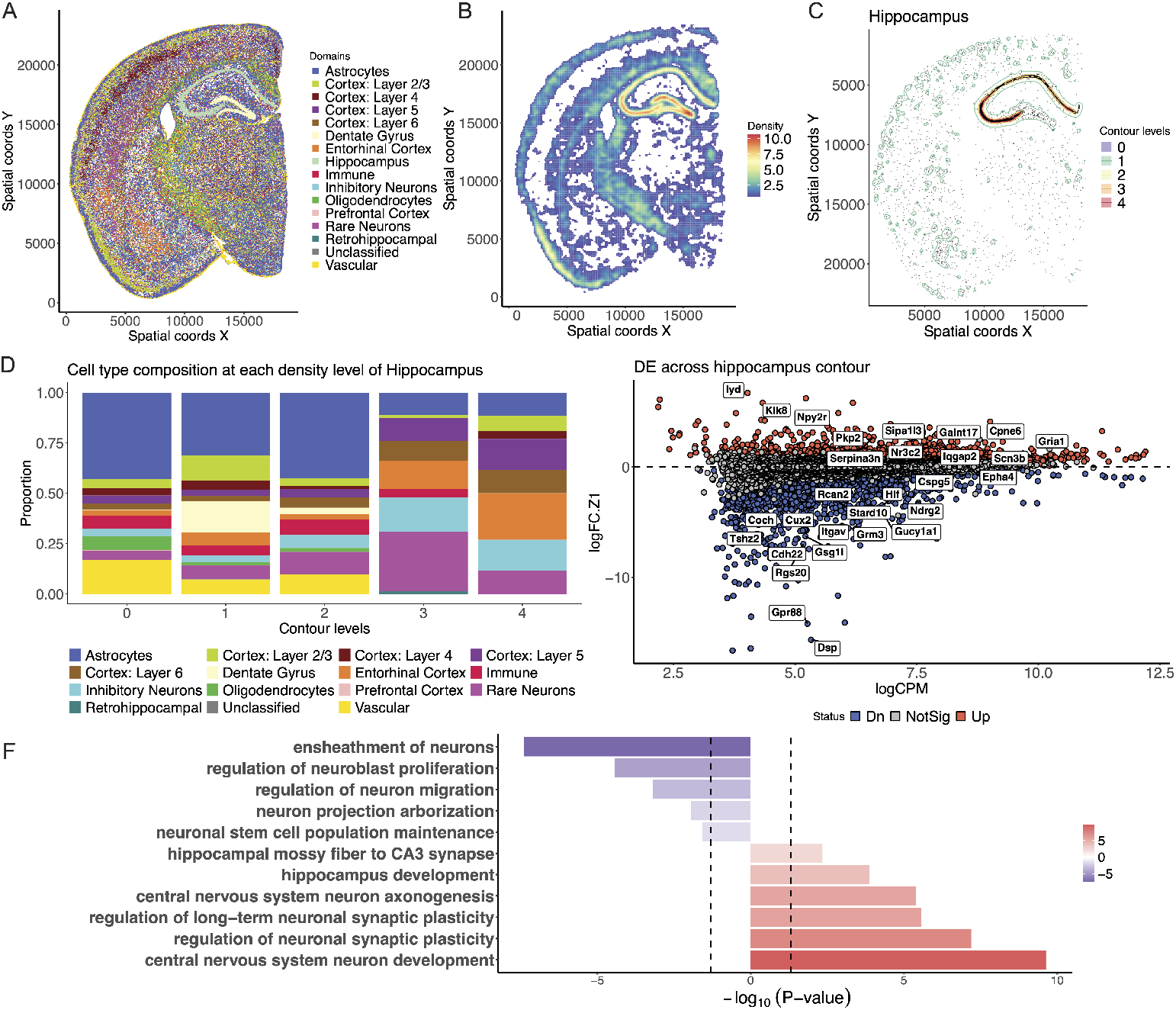
Spatial density analysis of Visium HD mouse brain data via scider. (A) Spatial plot with cell type annotation. (B) Heatmap of spatial density of cortex layer 2/3 and 6, dentate gyrus, hippocampus, immune infiltrates and oligodendrocytes. Grids of densities less than the median are filtered out in this visualization. (C) Spatial plot of hippocampus cells, overlaid by contour lines calculated from the spatial density of hippocampus. (D) Cell type composition at each contour level of hippocampus density cross the whole slide. (E) MA plots for gene expression changes associated with hippocampus densities. (F) Top neuro-associated genesets enriched in the differential expressed genes in E.

Cell-type deconvolution across these contours showed a monotonic rise in entorhinal-cortex excitatory neurons and inhibitory neurons as hippocampal density increased (Figure 6D). This is fully consistent with anatomical knowledge that the entorhinal cortex forms the major cortical input to the hippocampus and jointly supports episodic memory [31]. Inhibitory neurons, specifically GABAergic interneurons, also play an major role in hippocampal functions such as learning and memory formation [32].

We then aggregated UMIs within each contour to create pseudo-bulk profiles and conducted DE analysis with contour level as an ordered factor (see Methods). This analysis identified 1,395 genes that were significantly upregulated and 1,697 genes that were downregulated from level 0 to level 4 (FDR*<*0.05; Figure 4E). Among the top upregulated genes were GRIA1, PKP2 and KLK8, all canonical drivers of hippocampal synaptic plasticity, whereas strongly downregulated genes included CUX2 (cortical projection-neuron marker) and NDRG2 (astrocytic stress-response gene), mirroring the shift from neocortical and glial territories to mature hippocampal grey matter.

GO enrichment analysis of the DE genes was consistent with this interpretation (Figure 6F). Pathways linked to regulation of neuronal synaptic plasticity, central nervous-system neuron development and axonogenesis, and hippocampus development were positively enriched in hippocampus-dense regions, while terms associated with regulation of neruoblast proliferation and ensheathment of neurons were negatively enriched. Together, these findings demonstrate that scider not only recovers the expected hippocampal micro-anatomy but also captures the transcriptional trajectory from proliferative, myelinated cortical regions towards fully differentiated, plasticity-ready hippocampal circuits.

## Discussion

This article introduces the scider software for analyzing spatial transcriptomics data. Given cell type annotations, scider identifies spatial domains for downstream analyses based on the spatial distribution of specified cell type(s) of interest (COI). For the selected COI, scider estimates its spatial density by applying kernel density estimation (KDE) on a rectangular or hexagonal grid. Based on the spatial density estimation, scider provides two ways to define spatial domains, that is, regions of interest (ROIs) and contour regions.

ROIs are composed of spatially connected groups of adjacent grids derived from the KDE. Grid-level densities are used to construct a graph, where grids represent nodes and edge weights are the mean COIs densities between two adjacent grids. Community detection algorithms are applied to this graph and each community of nodes is defined as an ROI. ROIs provide a streamlined framework to construct cell type-oriented spatial domains for further analyses. Compared to pathologist-annotated ROIs, cell type-based ROI detection by scider is fully automated, reproducible and scalable. We have demonstrated that these ROIs can easily be annotated by gene expression data, which is not only less laborious but also enables the exploration of additional regions beyond those manually annotated by pathologists.

ROIs are particularly useful for large-scale spatial transcriptomics datasets, where manual annotation becomes impractical. ROIs enable the joint analysis of biological replicates to support more powerful and consistent identification of spatial patterns across samples. Reliable multi-sample analysis is critical for comparative studies across groups or conditions to support the identification of spatially resolved markers and molecular signatures of complex diseases. For example, we have shown that by jointly analyzing the two Xenium *in situ* replicates of the breast carcinoma sample, scider was able to identify a fourth cluster of ROIs that was not present in the individual analyses. Subsequent cell type composition, differential expression and pathway analyses helped elucidate distinct cellular profiles between the two DCIS#2 subtypes, highlighting spatial heterogeneity within DCIS#2-enriched regions. Although no clear batch effect was observed between these two replicates, we acknowledge that batch effects may present in other datasets, and normalization remains a challenge in multi-sample spatial data analysis. However, this ROI-based approach offers a practical framework for detecting and assessing potential batch effects.

The ROI-based approach has valuable applications in cancer biology, such as investigating tumor heterogeneity. We have demonstrated how our ROI-based analysis revealed molecular diversity within areas that pathologists had classified as DCIS#2 (Figure 3). Additionally, as noted in this study [11], not all DCIS lesions progress to invasive cancer. Distinguishing the molecular differences between DCIS subtypes is therefore critical for understanding disease progression. To this end, we also performed DE analysis to compare DCIS#1and DCIS#2-enriched regions (Figure 3A; cluster 1 versus clusters 2 and 4) in the Xenium *in situ* breast carcinoma replicates. COI cells, that is, DCIS and invasive tumor cells, were aggregated within each ROI to create pseudo-bulk samples. To account for the heterogeneity within pathologist-annotated DCIS#2 regions, a contrast was built to compare cluster 1 (DCIS#1) with the average of clusters 2 and 4 (DCIS#2). Supplementary Figure 4C and D summarize the DE results. Cluster 1 exhibited upregulation of genes associated with B cell function, lipid metabolism, and proliferation (e.g., MZB1, FASN, TFAP2A) and downregulation of genes linked to motility and immune signaling (e.g., ENAH, EGFR, CXCL16). GO analysis corroborated these findings, where we saw enrichment of kinase and metabolic processes and suppression of adaptive immune responses in cluster 1. Notably, the top 1 upregulated gene in cluster 1 was MZB1, which was reported to be an exclusive marker of DCIS#1 [11]. In addition, ROIs also serve as a systematic approach for cell type colocalization analysis. We have shown that cell type colocalization patterns vary significantly across different spatial regions, so that the ROI-based analysis reveals spatial patterns that a whole-slide approach may obscure [12].

Contour regions, on the other hand, define spatially contiguous areas with the similar COIs density. Contour regions enable the identification of spatial patterns associated with continuous variations in COIs density, providing a complementary approach to the discrete spatial regions defined by ROIs. In the first case study, we have used contour regions to investigate how cell type composition changed as tumor cell density increased. Consistent with what was observed in cell type colocalization analysis, cell type composition also displayed distinct patterns across spatial regions. For example, when analyzing the entire invasive tumor-enriched region, we observed similar T cell proportions across all contour levels (Figure 4E). However, within ROIs 7 and 17, T cell proportion decreased with increasing invasive tumor density (Supplementary Figure 9). This suggests that T cell infiltration may be more pronounced in certain tumor regions, which was masked in a whole-slide analysis.

Furthermore, contour-based DE analysis identifies microenvironment-specific cell states and transcriptional trajectories linked to increasing COIs density. In the Xenium case study, we demonstrated how this approach helped dissect intratumor heterogeneity. However, the contour-based DE analysis is potentially susceptible to transcript diffusion. This is especially true in regions with high cellular density or dominated by specific cell types, where transcripts from the COIs may diffuse into neighboring cells and confound the DE results. For example, when comparing B cells across invasive tumor contours, we observed several upregulated genes, including KRT7, EPCAM and KRT8, which are typically viewed as epithelial or tumor cell markers (Figure 5E, top). Such a result may also arise from inaccuracies in cell segmentation or cell type annotation. We anticipate that as upstream cell segmentation and annotation methods improve, the accuracy of contour-based DE analysis, as well as other scider analyses, will also improve.

In the meantime, these considerations led us to adopt the pseudo-bulking approach for DE analysis presented in this aricle. By aggregating cells within each ROI or contour region, pseudo-bulking helps mitigate the impact of transcript diffusion. This strategy also enables seamless integration with well-established tools such as limma and edgeR for downstream analyses, including gene set enrichment and pathway analysis. Both limma and edgeR provide rigorous statistical frameworks that account for biological variation across replicate samples, which is essential when analysing multi-sample spatial transcriptomics data.

For imaging-based spatial transcriptomics platforms like Xenium and MERSCOPE, the scider pipeline can be extended to a cell-free approach to further reduce technical artifacts arising from cell segmentation and cell type annotation ambiguity. By aggregating observed gene molecule counts within spatial grids, scider can construct expression profiles at the grid-level, similar to the Visium and Visium HD platforms but with a smaller gene panel, while still retaining some single-cell resolution within each grid. In this framework, KDE can be applied to an individual gene or gene-set, rather than at the cell-type level, minimizing its dependence on the upstream cell-segmentation and cell-annotation processes. For example, scider allows the identification of gene-specific ROIs by performing KDE on cell coordinates weighted by the expression level of the gene(s) of interest. As density estimation is based on cell locations, the underlying tissue organization is preserved. Supplementary Figure 11 illustrates the spatial density of ACTA2 and KRT15 expression in the Xenium breast carcinoma dataset, both myoepithelial cell markers used in the original study [11]. We showed that ROIs identified for elevated ACTA2 or KRT15 expression largely overlap with regions enriched with myoepithelial cells (Supplementary Figure 1E, left). Similarly, ROIs were identified for elevated expression of T cell markers as shown in Supplementary Figure 12, which are primarily concordant with T cell-enriched regions as shown in Supplementary Figure 1F (right). Furthermore, building on this approach, scider can be extended to support cell-cell communication analysis by examining the kernel density distributions of ligands and receptors to identify spatial hotspots where paired ligand-receptor signals colocalize.

In addition, we have also demonstrated that scider can be applied to sequencing-based spatial transcriptomics platforms such as 10x Visium and Visium HD. In our analysis of the Visium HD mouse brain data, each 8 × 8 *μ*m grid was treated as a single cell, as Visium HD does not provide true cell segmentation. However, each grid may capture transcripts from multiple cells or cell fragments. This limits the accuracy of cell type-specific analyses, and results can be sensitive to grid size and resolution. Large grids may lead to mixed signals from different cell types and obscure spatial patterns, while small grids may result in a lower signal-to-noise ratio.

In summary, scider introduces a novel and streamlined framework for spatial transcriptomics analysis. It defines spatial domains based on the spatial distribution of a specified COI, represented as ROIs or contour regions. KDE is used to estimate the spatial density of the COI and can be flexibly applied to heterogeneous types of tissues and technologies through user-supplied parameters. scider supports a range of downstream analyses including cell type colocalization, cell type composition and DE analysis, along with efficient and flexible visualization options for data exploration and interpretation. scider is publicly available as an R package on Bioconductor (https://bioconductor.org/packages/scider/).

## Methods

### scider workflow

#### Spatial density estimation

Briefly, kernel density estimation (KDE) estimates the spatial density by spreading the probability mass of each data point into its surrounding neighborhood using a kernel function. scider takes a SpatialExperiment object as input, which contains the spatial coordinates of each cell. For a given cell type of interest (COI), scider calculates KDE based on the spatial locations of all COIs, using either a hexagonal or rectangular grid across the tissue. The density estimate is scaled so that its value represents the expected number of cells on each grid. The `gridDensity`function of scider implements the KDE, internally leveraging the spatstat R package for rectangular grids and the hexDensity R package for hexagonal grids.

By default, KDE is carried out using a grid size of 100 *μ*m. In practice, we recommend adjusting this value based on the characteristics of the tissue and the technology used. The degree of smoothing in KDE is controlled by the ‘bandwidth’ parameter, which scider automatically selects using the crossvalidated selector from the spatstat R package [33], though it can also be manually supplied. Edge correction is applied to account for the underestimation of density near tissue sample boundaries.

#### ROI detection

Spatial density estimated for the cell types of interest (COIs) is used to construct a weighted and undirected graph, where each grid is a node and adjacency defines edges. If two grids are adjacent to each other, an undirected edge exists between the two grids. Each edge is weighted by the average density of the COIs between the two grids it connects. The graph is then used to identify regions of interest (ROIs) by community detection, where grids of each community comprise an ROI. ROI detection is implemented in scider as the `findROI`function, where network analyses are performed using the igraph R package [34].

#### Cell type colocalization

Cell type colocalization describes the spatial distribution of two or more cell types in a given region. Importantly, conclusions about colocalization are region-specific and may vary across different areas, even within the same sample.

In scider, we use the correlation of spatial densities between two cell types to measure their colocalization. Specifically, the Pearson correlation coefficient is computed using the spatial densities of the two cell types across all grids within a specified region. The correlation can be calculated using grids across the whole slide or within each ROI. Significance of the correlation is tested by a modified t-test to account for spatial autocorrelation [35, 36].

Additionally, cell type colocalization can be jointly assessed across multiple ROIs. The average correlation coefficient is calculated to summarize the overall correlation, and Fisher’s method [37] is used to combine the p-values across all ROIs within a tissue.

Cell type colocalization analysis is implemented in scider as the `corDensity` function.

#### Contour detection and cell type composition analysis

Contour detection is performed given the spatial density estimation of COIs, with contours defined as the boundaries of regions at a given density level. Contours are calculated using the `getContour` function in scider, based on the `geom_contour` function in ggplot2 [38].

Cell type composition analysis can be performed at each contour level by allocating each cell to the contour level it resides. Specifically, the sf R package [39] is used to carry out the spatial operations. Contour isolines are first converted into polygons. The area corresponding to each contour level is defined by computing the difference between polygons of adjacent levels. Each cell is then allocated to the corresponding contour level based on the spatial region it falls within. Contour level allocation is implemented in scider as the `allocateCells`function, by which an extra column is added to the colData of the SpatialExperiment object to store the contour level for each cell.

At each contour level, the proportion of each cell type is calculated by dividing the number of cells of that type by the total number of cells at that contour level. The cell type composition can be visualized in scider using the `plotCellCompo`function.

#### DE analysis

For a given tissue sample, differential expression (DE) analysis can be performed at the ROI level or the contour level, or even both. The pseudo-bulk strategy is generally recommended for this type of DE analysis. Briefly, pseudo-bulking is performed by aggregating cells within each ROI or at each contour level using the `spe2PB`function to create a DGEList object, a data object used in the edgeR package, containing the pseudo-bulk expression values. The `spe2PB`function also supports separating cell types during pseudo-bulking, which is useful for cell type-specific DE analysis. DE analysis can then be performed using the quasi-likelihood approach in edgeR v4.0 [40] or the `voomLmFit`pipeline [41]. This makes scider fully compatible with all existing limma and edgeR functionalities, allowing any arbitrarily complex experimental design and a wide range of downstream tasks including KEGG pathway and gene ontology (GO) analysis. More importantly, the flexibility of these frameworks allows DE analysis to be extended to multi-sample datasets, where modeling sample effects and biological variation is highly essential.

### Analysis of Xenium in situ data

#### Data preprocessing

The Xenium *in situ* breast carcinoma data were obtained from a published study [11]. In this case study, we focused on the two Xenium *in situ* replicates of Sample 1. Cell type labels were sourced from the same study, with subgroups belonging to broad cell type categories were merged. In particular, we merged the following cell types: ACTA2^+^ myoepithelial cells and KRT15^+^ myoepithelial cells were merged into “myoepithelial cells”; CD4^+^ T cells and CD8^+^ T cells were merged into “T cells”; DCIS#1 and DCIS#2 were merged into “DCIS” unless otherwise specified (e.g., in Figure 3C); macrophages 1 and macrophages 2 were merged into “macrophages”; and LAMP3^+^ dentritic cells and IRF7^+^ dentritic cells were merged into “dentritic cells”. Spatial plots with the updated cell type annotation are available in Supplementary Figures 1, 3A and 3B.

All analyses on this Xenium dataset were performed in R v4.5.1 with scider v1.5.6.

#### ROI-based analysis

For ROI-based analyses, the cell type of interest (COI) was chosen to be the combination of ductal carcinoma *in situ* (DCIS) and invasive tumor cells. Histology/pathology and scFFPE-seq guided ROIs were obtained from the same study and used to compare with scider-detected ROIs. For simplicity, we had referred to these histology/pathology and scFFPE-seq guided ROIs as “pathologist-annotated ROIs” throughout this paper.

For both replicates, cell type densities were calculated based on 50 *μ*m hexagonal bins. Bandwidth for KDE was set to be half the width of each hexagon (25 *μ*m). We chose a smaller bandwidth (compared to the contour-based analysis) to achieve a finer spatial resolution in detected ROIs at the given grid size. Supplementary Figure 2 shows ROI detection results obtained with varying bandwidth values. Heatmaps of the estimated spatial density of COI for both replicates are shown in Supplementary Figure 3C and D. Grids with low COI densities (less than 0.5) were filtered out before ROI identification. Method for community detection was by default set to “greedy” in the `findROI`function, using the greedy modularity optimization algorithm [42] via the cluster fast greedy function from the igraph R package [34]. ROIs of size less than 20 grids were filtered out.

ROIs were then used to create pseudo-bulk samples by aggregating COI cells within each ROI. The `spe2PB`function was used to create a DGEList object containing the pseudo-bulk expression values. The pseudo-bulk samples were then used to perform multi-dimensional scaling (MDS) analysis using the `plotMDS`function from limma v3.64.1 [43]. Top 50 genes with the largest log2-fold-changes between every two samples were used to generate the MDS plot. In the analysis of replicate 1, k-means clustering of the ROIs was sensitive to random seed initialization and produced unstable results with *k* set to either 3 or 4. We therefore identified the clusters by visual inspection of the MDS plot (Figure 2E) and that resulted in 4 clusters. In the joint analysis of both replicates, ROI clusters were stably identified by k-means clustering on the MDS coordinates with *k* set to 4 (Figure 3A). Cell type composition of each ROI was calculated by dividing the number of cells of that type by the total number of cells in that ROI, with (Figure 3C, left) or without (Figure 3C, right) the COI itself included.

We first performed DE analyses to compare COI cells between ROIs of different clusters (Supplementary Figure 4). Pseudo-bulk samples were created by aggregating the COI cells within each ROI. The replicate effect was accounted for by including replicate ID as a factor in the design matrix.

We also performed cell type-specific analyses to compare several non-COI cell types in ROIs of cluster 2 versus those of cluster 4, both DCIS#2-enriched. Pseudo-bulk samples were created by aggregating cells within each ROI for every cell type. This analysis was performed for endothelial cells, myoepithelial cells, macrophages, stromal cells, and T cells, while adjusting for the replicate effect (Figure 3E).

#### Contour-based analysis

Contour-based analyses were next performed on replicate 1, where invasive tumor cells were selected as COI. Cell type densities were calculated based on 50 *μ*m hexagonal bins. Bandwidth for KDE was also set to 50 *μ*m. Cell type colocalization was performed on the whole slide (Figure 4D), in each ROI (Figure 4C, Supplementary Figures 7 and 8), and across all ROIs (Figure 4B). Cell type composition was computed by dividing the number of cells of each cell type by the total number of cells at each contour level. Results were visualized with the COI excluded, both across all ROIs (Figure 4E) and for each individual ROI (Figure 4F and Supplementary Figure 9).

We then performed DE analysis to compare cells at different contour levels of the COI. By default, getContour generates contours at evenly spaced contour levels, given the total number of levels defined by users. Alternatively, contours can be manually defined using user-specified density cutoffs. For interpretability, we generated contours in this analysis by explicitly setting the density breakpoints at 2, 4, 6, 8 and 10, yielding six contour levels. That is, level 0 corresponds to grids with COI density less than 2, level 1 to grids with COI density between 2 and 4, and so on. The density value represents the expected number of COI cells within each grid.

Pseudo-bulk samples were then created by aggregating cells at each contour level for each cell type. The `spe2PB`function was used to create a DGEList object, with each sample representing a specific cell type at a defined COI density level. For each cell type (invasive tumor, stromal and B cells), a design matrix was constructed using contour level breaks as the covariate, which corresponds to the minimum COI density at each contour level.

#### DE and pathway analysis

All pseudo-bulk DE analyses on this Xenium *in situ* dataset were performed using the `voomLmFit`pipeline [41] in edgeR v4.6.2. The log-count per million (logCPM) values of the pseudo-bulk profiles were normalized using the cyclic loess normalization [44], followed by the downstream DE analysis using `voomLmFit`with sample weights. A false discovery rate (FDR) threshold of 5% was used to define differential expression. KEGG pathway analysis was performed using the `kegga`function, and GO analysis using the `goana`function, both from limma [43].

### Visium HD data analysis

#### Data preprocessing

The 10x Visium HD mouse brain dataset was obtained from the official 10x Genomics repository (https://www.10xgenomics.com/datasets/visium-hd-cytassist-gene-expression-libraries-of-mouse-brainhe). Count matrices of 8 *μ*m bin size were used. Bin-level quality control metrics, including total UMI count, number of detected genes, and percentage of mitochondrial transcripts, were calculated using the scuttle R package. Bins with fewer than 30 total counts, fewer than 10 detected genes, or more than 20% mitochondrial content were excluded. After filtering, expression values were normalized using the deconvolution-based size factor estimation from scran.

#### Cell type annotation

Bin-level normalized profiles were annotated using SingleR [45] by comparing each bin’s transcriptomics profile against the Allen Mouse Brain single-cell data [46]. SingleR employs a Wilcoxon rank-sum test to identify reference marker genes and assigns each bin the label with the highest correlation to cell type from the reference profiles. To ensure robust calls, only bins whose top label score was ≥ 0.1 and whose difference (delta) to the next-best score was ≥ 0.05 were retained; all others were excluded from downstream analyses.

Assigned labels were then aggregated into broader anatomical domains: vascular-associated labels (VLMC, endothelial, pericytes) as “Vascular”; microglia and perivascular macrophages as “Immune”; classical glial types as “Astrocytes” or “Oligodendrocytes”; excitatory neuron subtypes into cortical layers (Layers 2/3, 4, 5, 6) or entorhinal cortex; hippocampal neurons as “Dentate Gyrus” or “Cornu Ammonis/Subiculum”; and inhibitory interneurons by lineage (VIP^+^, Pvalb^+^, Sst^+^). Rare or lowconfidence labels were marked “Unclassified”. These domain annotations provided the framework for subsequent density estimation and contour analyses.

#### DE and GO analysis

DE analysis on the Visium HD dataset was carried out as described in the previous section. TMM normalization was applied to logCPM values [47]. Genes with FDR *<* 0.05 were selected as significant differential expressed genes. GO analysis was conducted using the `goana`function.

## Data and code availability

Both Xenium and Visium HD data are publicly available on the 10x Genomics website; code used in this manuscript is available at https://github.com/ChenLaboratory/scider_manuscript_code.

## Acknowledgments

M.L. was supported by Melbourne Research Scholarship and CSL Translational Data Science Scholarship. N.L. was supported by the South Australian immunoGENomics Cancer Institute (SAiGENCI) initiation funds and Adelaide Centre of Epigenetics (ACE) initiation funds. Y.C. was supported by Medical Research Future Fund Investigator Grant (1176199). We thank the support provided by the Estate of the Judith Corrie Philpots, and by the Victorian State Government (Operational Infrastructure Support to WEHI).

We thank Lizhong Chen for valuable discussions regarding methods for colocalization analysis. We thank Jane Visvader and Michael Milevskiy for detailed feedback on the Xenium case study. We thank Saskia Freytag, Sarah Best, Rory Bowden and Givanna Putri for their feedback on the manuscript.

## Competing interests

All authors declare that they have no competing interests.

## Author contributions

M.L. conceptualized and developed methods, wrote the software, carried out analyses on the Xenium dataset, wrote and finalized the paper. N.L. co-developed and co-wrote the software, carried out analyses on the Visium HD dataset and co-wrote the paper. Q.H.N. developed the hexagonal kernel density estimation method and co-wrote the paper. Y.C. conceived and supervised the project, co-developed the software and co-wrote the paper. All authors contributed to and checked the final manuscript.

## Notes

### Competing Interest Statement

The authors have declared no competing interest.

