## Supplementary materials for "Preserving tissue structure through density-based spatial analysis with scider"

### Supplementary Figures

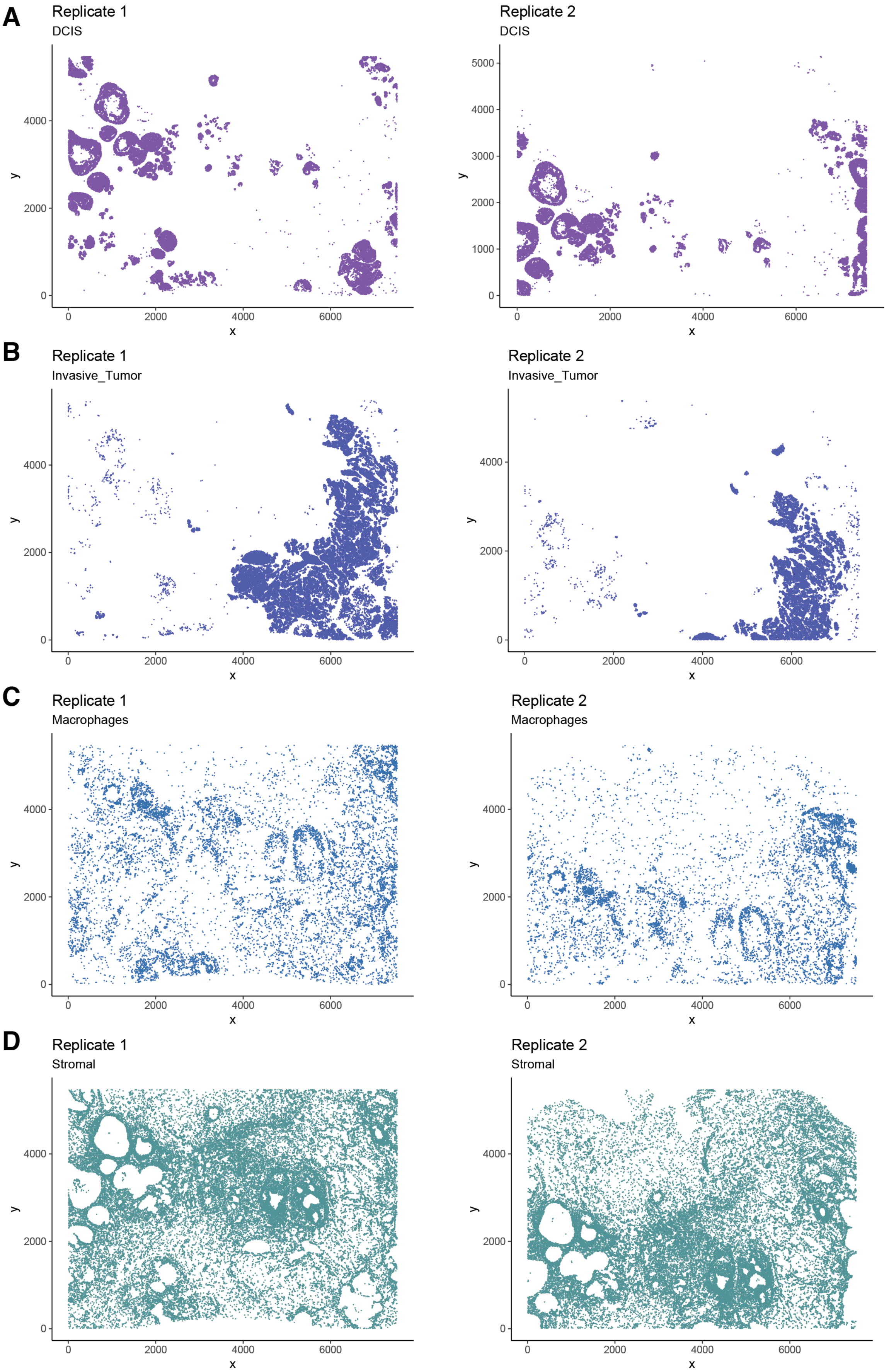


**Figure S1:** Spatial plots of each cell type in the Xenium breast carcinoma dataset for replicate 1 (left) and replicate 2 (right). Panels (A)–(D) are DCIS (DCIS #1 and DCIS #2 combined), invaive tumor, macropages and stromal cells, respectively.

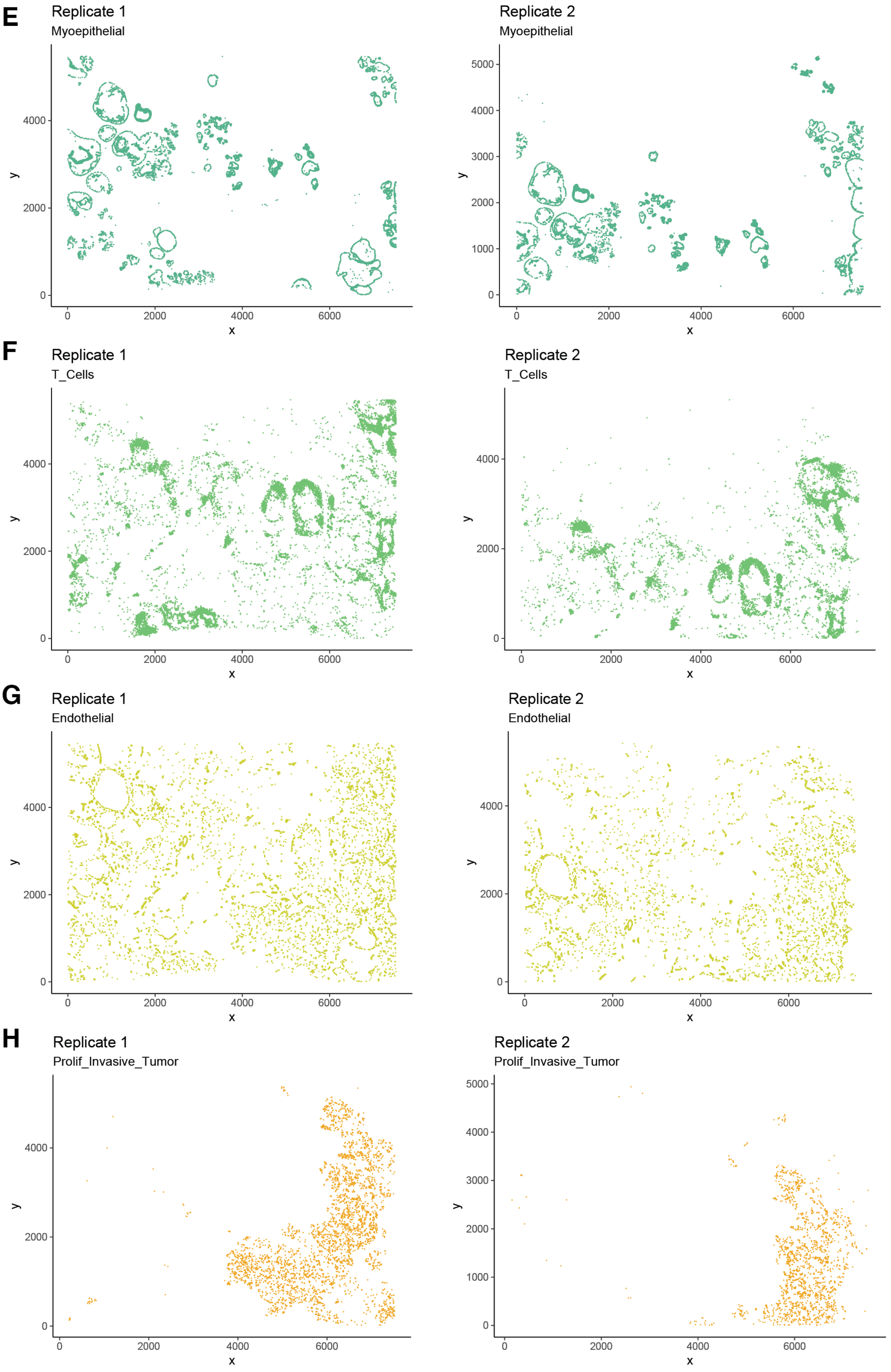


**Figure S1:** Spatial plots of each cell type in the Xenium breast carcinoma dataset for replicate 1 (left) and replicate 2 (right), continued. Panels (E)–(H) are myoepithelial, T cells, endothelial cells and proliferative invasive tumor, respectively.

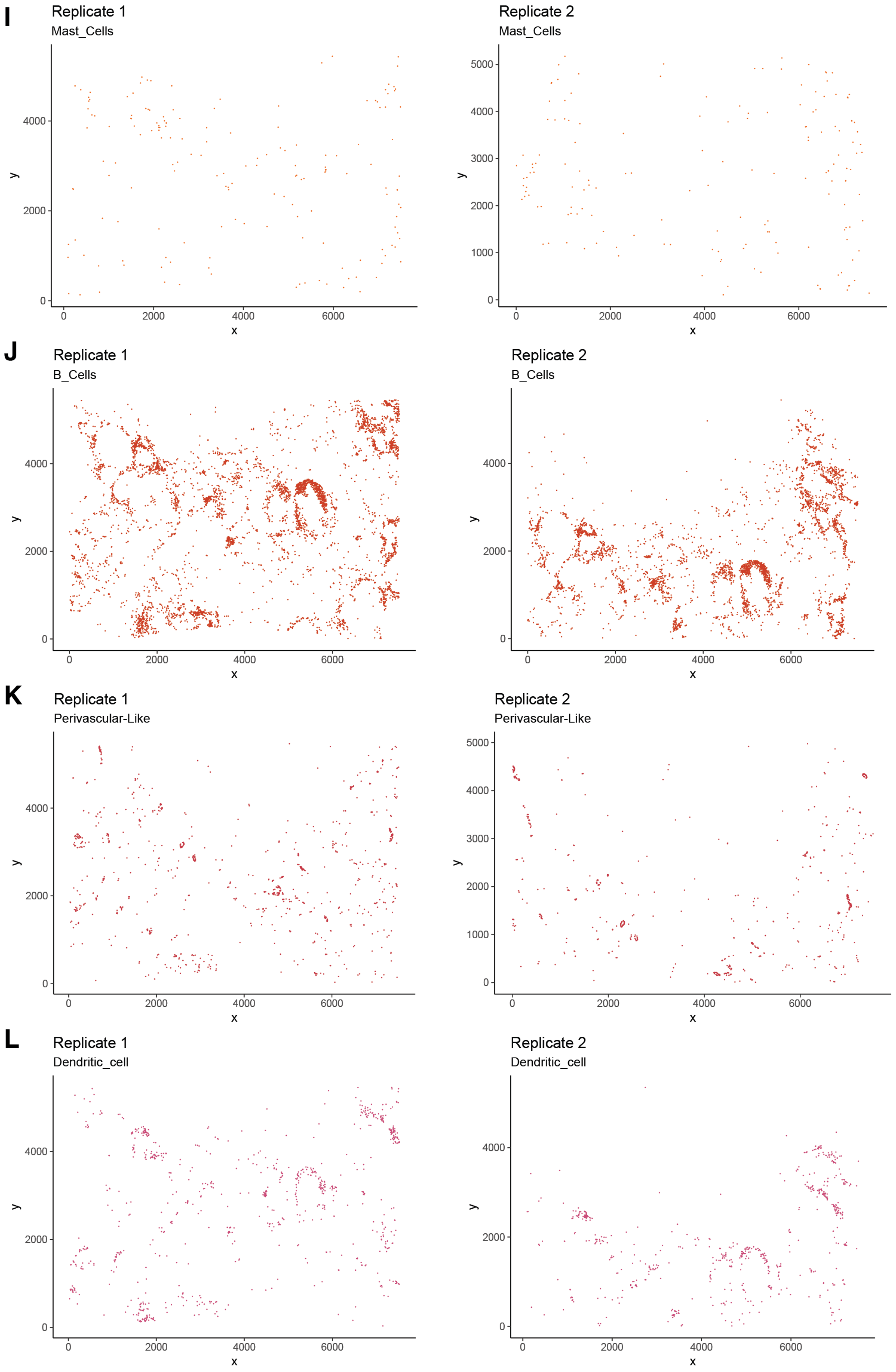


**Figure S1:** Spatial plots of each cell type in the Xenium breast carcinoma dataset for replicate 1 (left) and replicate 2 (right), continued. Panels (I)–(L) are mast cells, B cells, perivascular-like cells and dendritic cells, respectively.

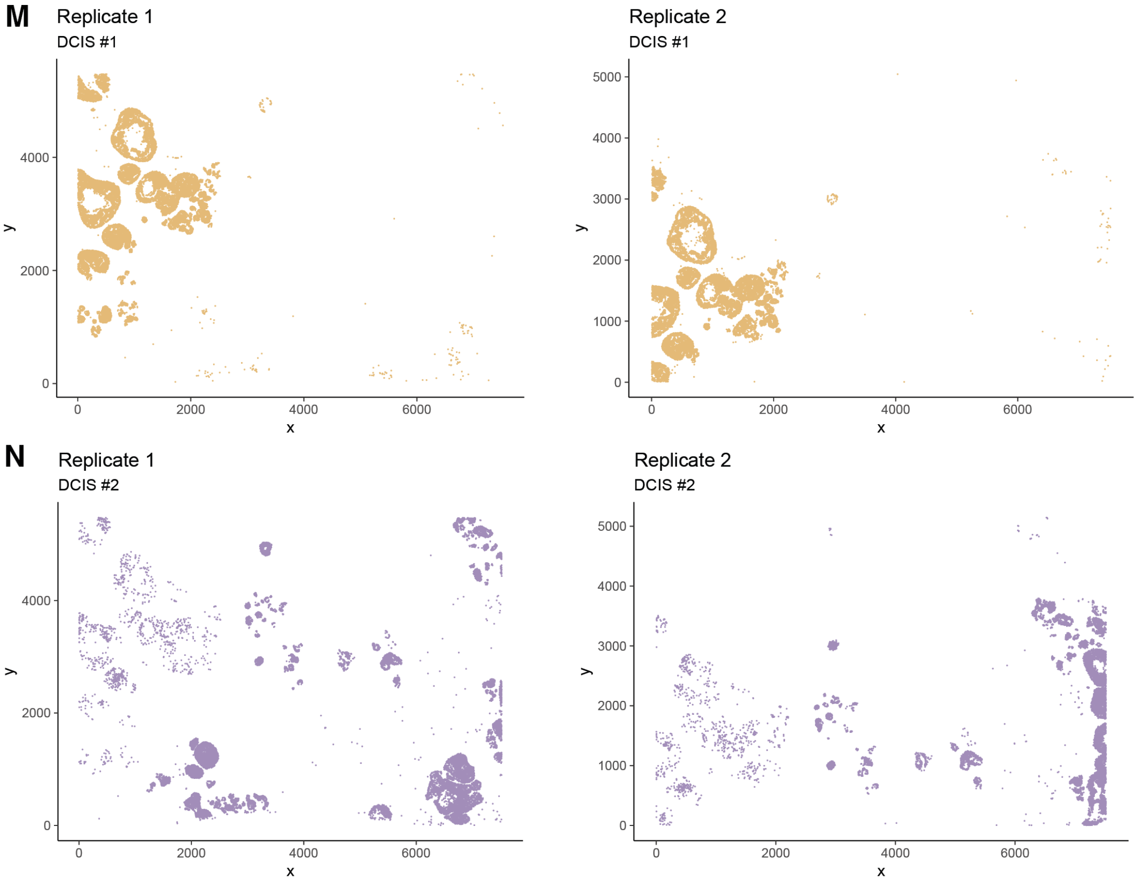


**Figure S1:** Spatial plots of each cell type in the Xenium breast carcinoma dataset for replicate 1 (left) and replicate 2 (right), continued. Panels (M) and (L) are DCIS #1 and DCIS #2, respectively.

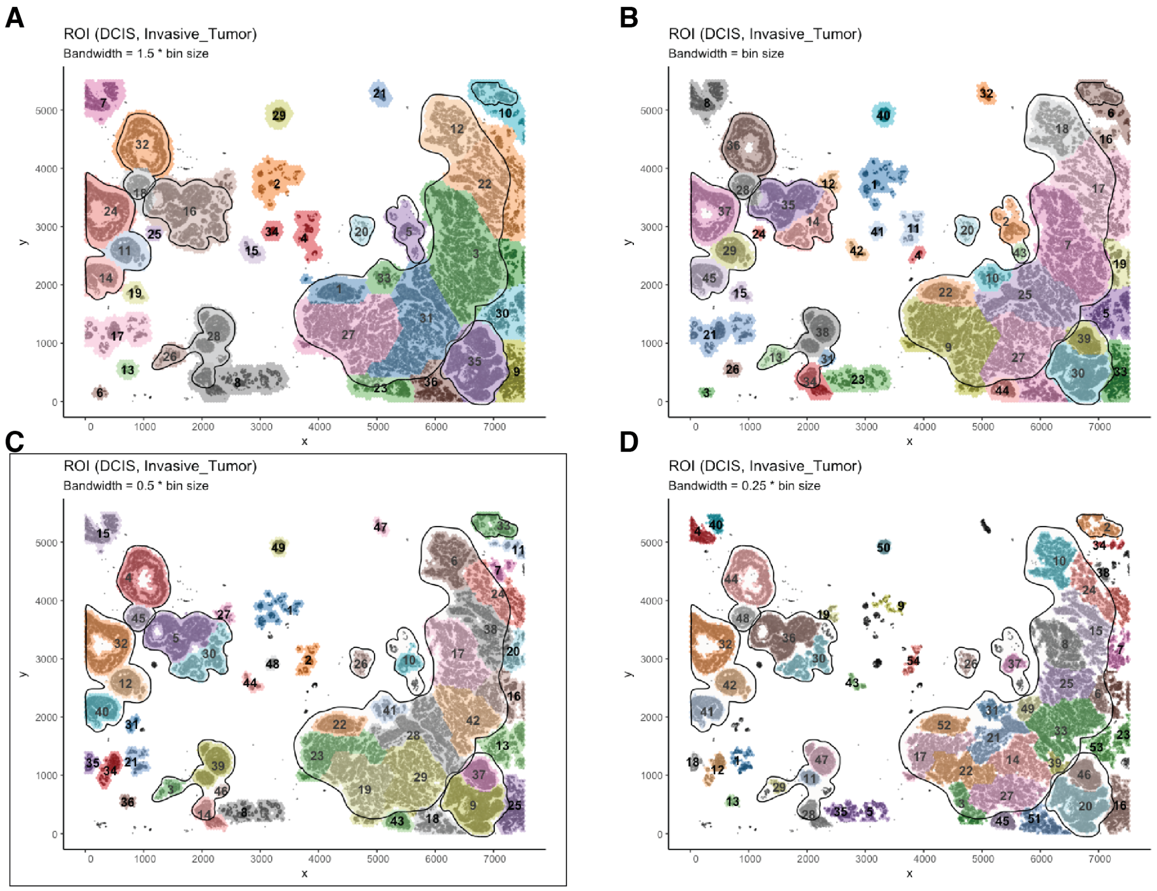


**Figure S2:** ROI detection on the Xenium breast carcinoma sample (replicate 1) with varying bandwidths. The cell type of interest (COI) was chosen to be the combination of ductal carcinoma in situ (DCIS) and invasive tumour cells. The spatial density of the COI was estimated by dividing the slide into 50$\mu$m-wide hexagonal grids for kernel density estimation (KDE). Grids with low COI densities (less than 0.5) were filtered out. A weighted and undirected graph was constructed based on the spatial density of the COI. Regions of interest (ROIs) were identified via community detection by the greedy modularity optimization algorithm. ROIs with less than 20 grids were filtered out. (A) ROI detection with a bandwidth of 75$\mu$m (1.5$\times$bin size). (B) ROI detection with a bandwidth of 50$\mu$m (the same as bin size). (C) ROI detection with a bandwidth of 25$\mu$m (0.5$\times$bin size). (D) ROI detection with a bandwidth of 12.5$\mu$m (0.25$\times$bin size).

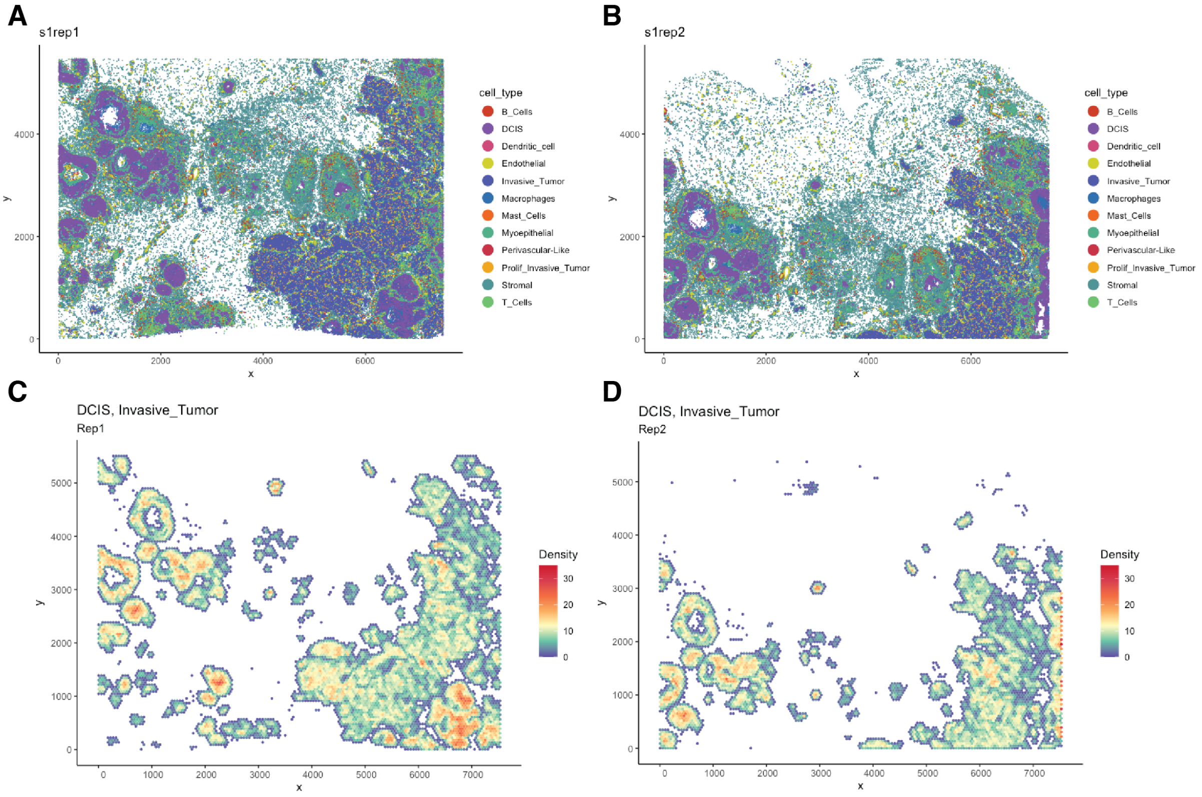


**Figure S3:** Overview of the two Xenium breast carcinoma replicates. (A) Spatial plot with cell type annotation in replicate 1. This is the same as Figure 2A in the main text. (B) Heatmap of spatial density of DCIS and invasive tumor cells in replicate 1. This is the same as Figure 2B in the main text. (C) Spatial plot with cell type annotation in replicate 2. (D) Heatmap of spatial density of DCIS and invasive tumor cells in replicate 2.

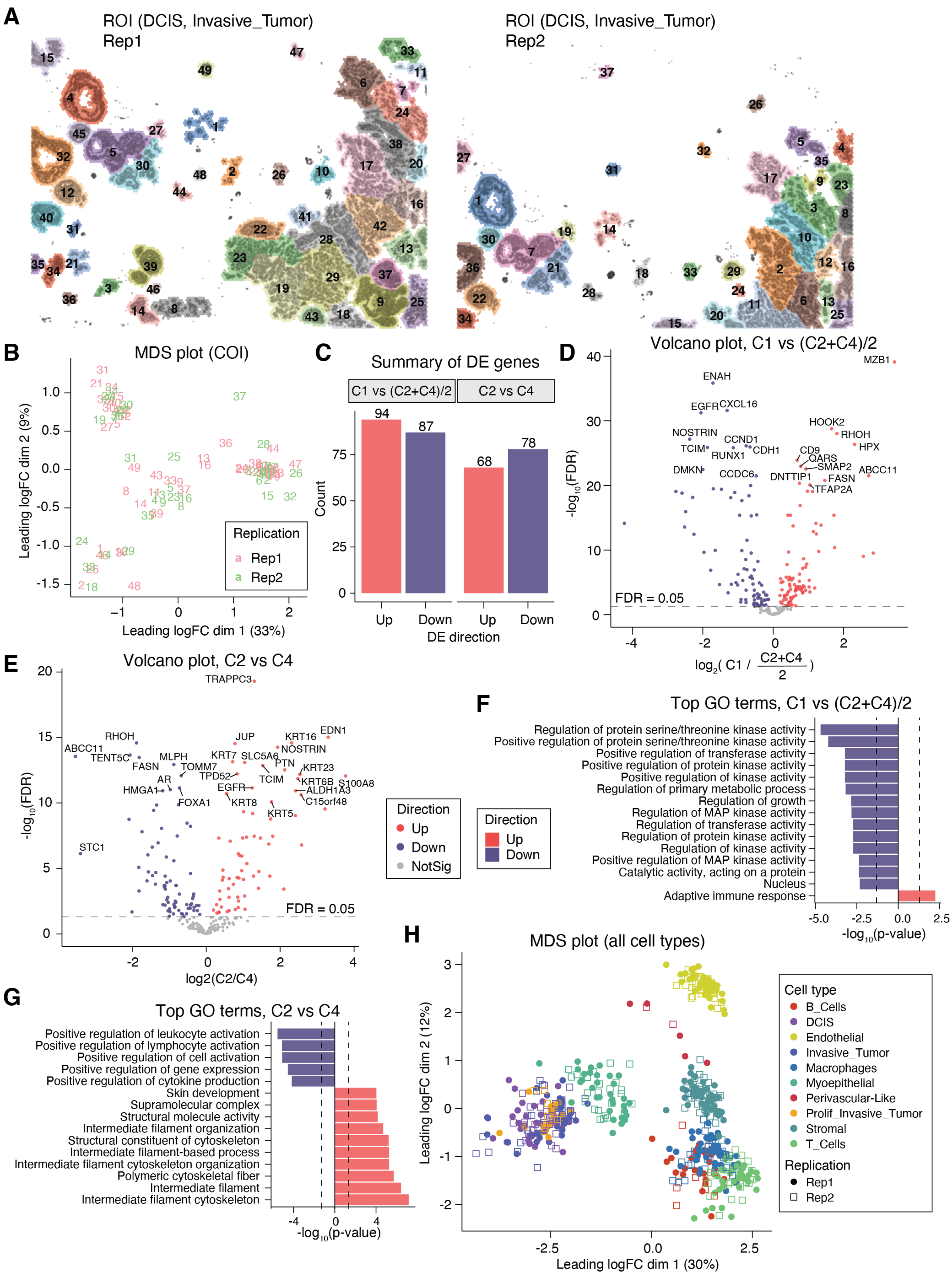

**Figure S4:** ROI-based joint analysis of the two Xenium breast carcinoma samples (replicate 1 and replicate 2). DCIS and invasive tumor cells are used as COI and highlighted in the background. (A) ROIs detected by scider on repliate 1 (left) and replicate 2 (right). The left panel is the same as Figure 2C in the main text. (B) MDS plot of gene expression in COI cells from ROIs in both replicates. Pseudo-bulk samples are created by aggregating COI cells within each ROI. This is the same as Figure 3A in the main text, but annotated by ROI IDs and replicates. (C) Summary of DE genes comparing COI cells in different ROI clusters identified in Figure 3A. (D) Volcano plot comparing COI cells from cluster 1 (C1) ROIs with those from cluster 2 (C2) and cluster 4 (C4). (E) Volcano plot comparing COI cells from C2 ROIs with those from C4. (F) Top GO terms comparing COI cells from C1 ROIs with those from C2 and C4. (G) Top GO terms comparing COI cells from C2 ROIs with those from C4. (H) MDS plot of gene expression in all cell types from ROIs in both replicates. Pseudo-bulk samples are created by aggregating cells within each ROI for every cell type. This is the same MDS plot as Figure 3E, annotated according to cell types and replicates.

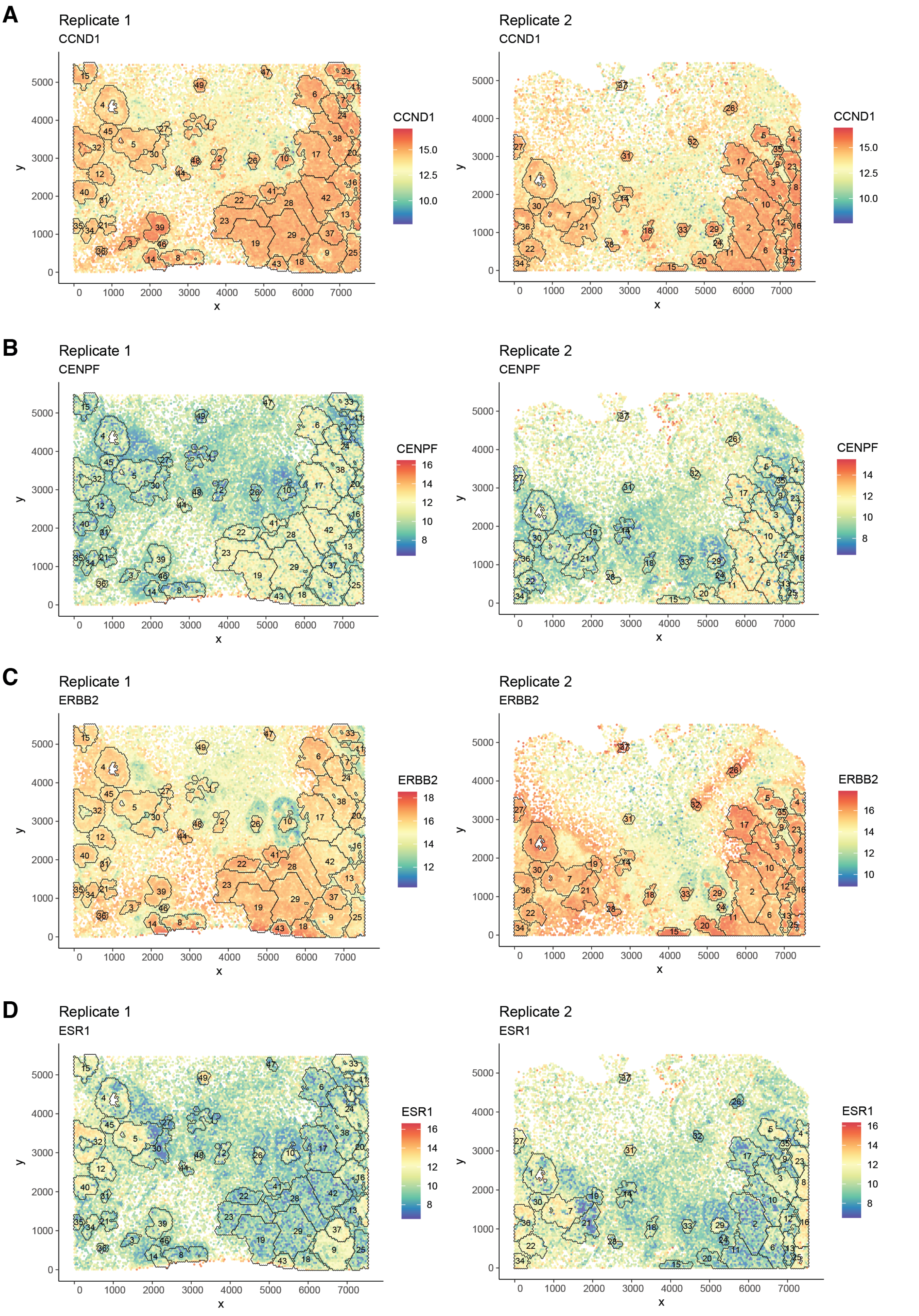


**Figure S5:** Spatial plot of marker gene expression overlaid by DCIS and invasive tumor ROIs in the Xenium breast carcinoma sample for replicate 1 (left) and replicate 2 (right). Panels (A)–(D) are CCND1, CENPF, ERBB2 and ESR1, respectively, all on the logCPM scale.

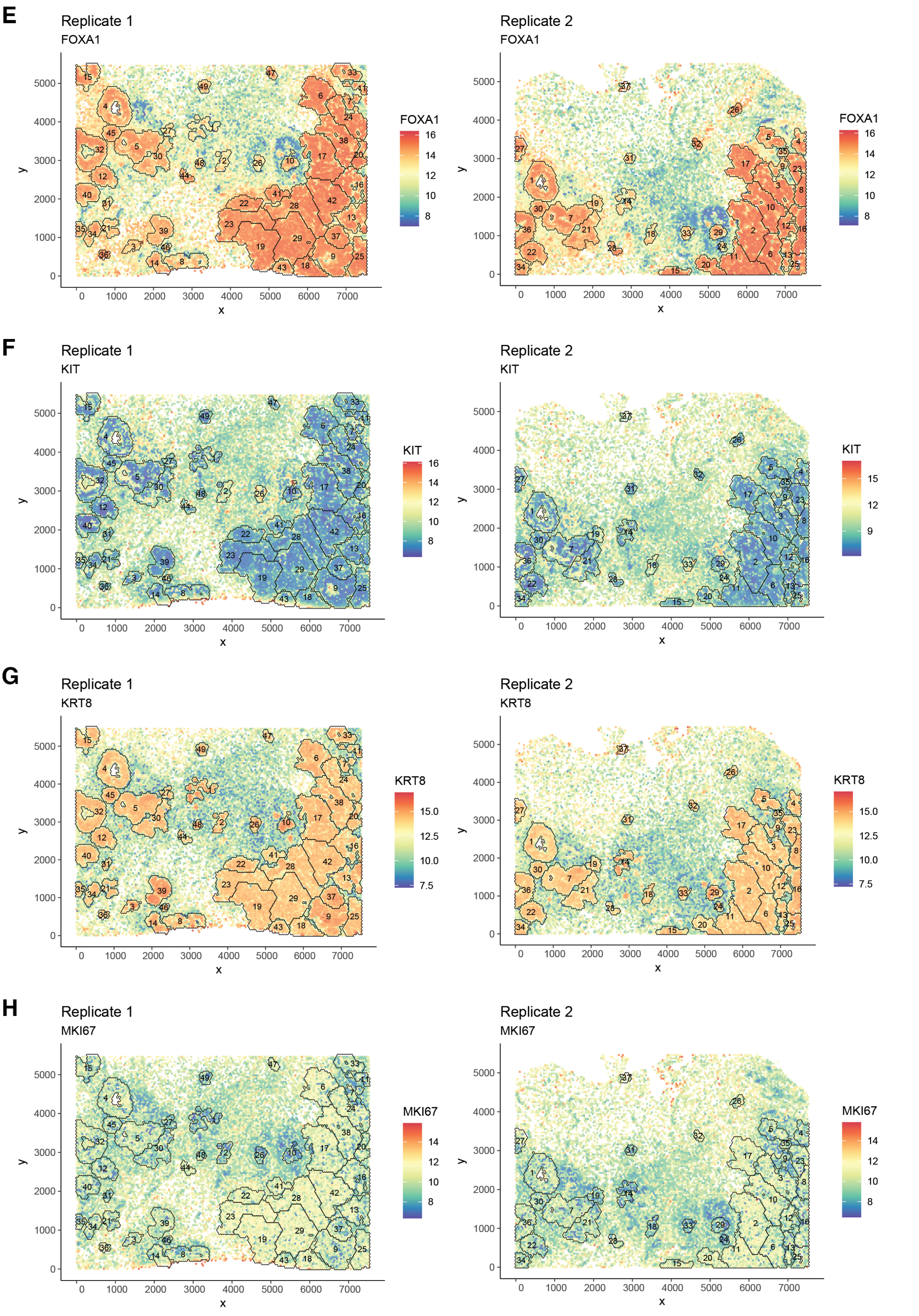


**Figure S5:** Spatial plot of marker gene expression overlaid by DCIS and invasive tumor ROIs in the Xenium breast carcinoma sample for replicate 1 (left) and replicate 2 (right), continued. Panels (E)–(H) are for FOXA1, KIT, KRT8 and MKI67, respectively, all on the logCPM scale.

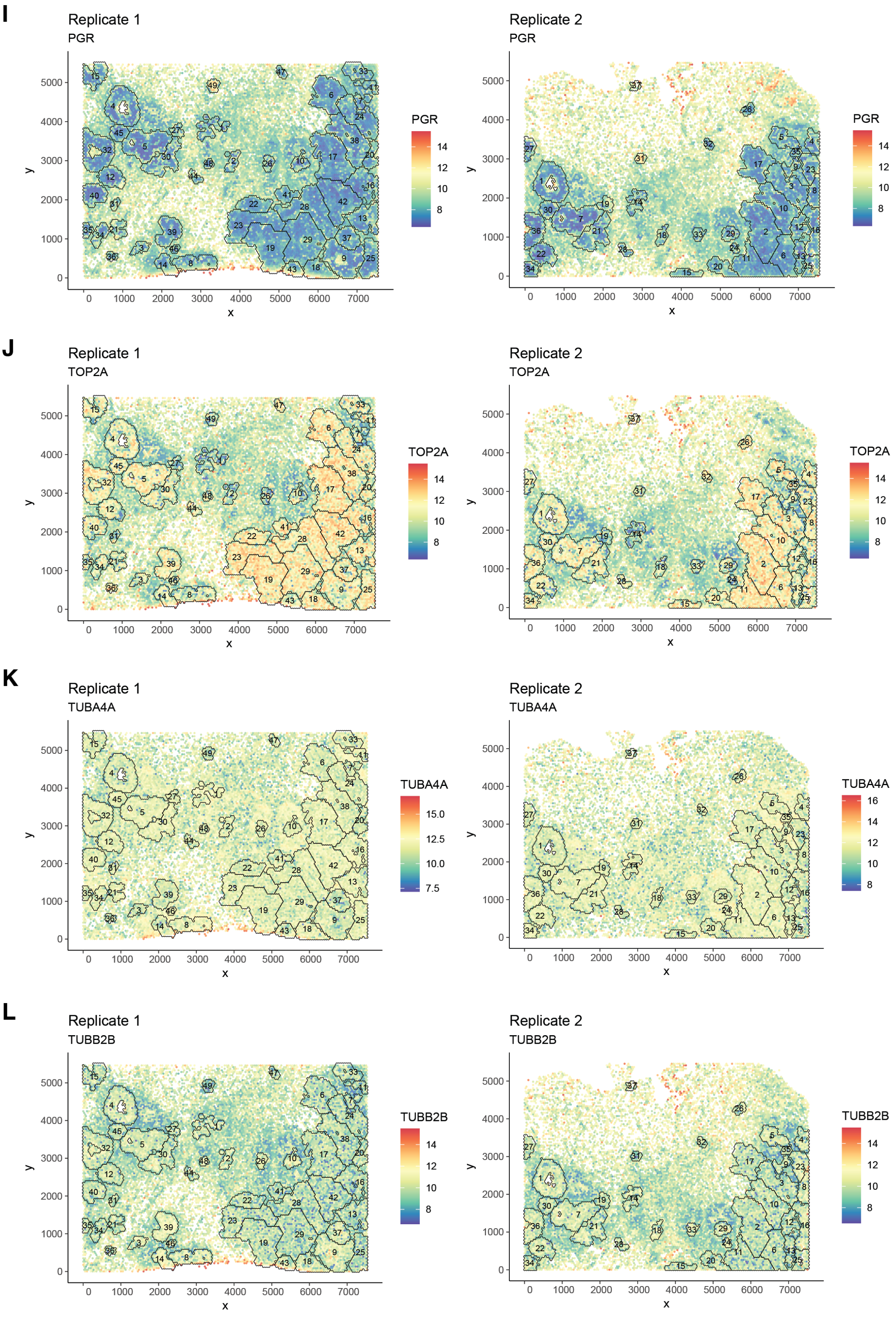


**Figure S5:** Spatial plot of marker gene expression overlaid by DCIS and invasive tumor ROIs in the Xenium breast carcinoma sample for replicate 1 (left) and replicate 2 (right), continued. Panels (I)–(L) are for PGR, TOP2A, TUBA4A and TUBB2B, respectively, all on the logCPM scale.

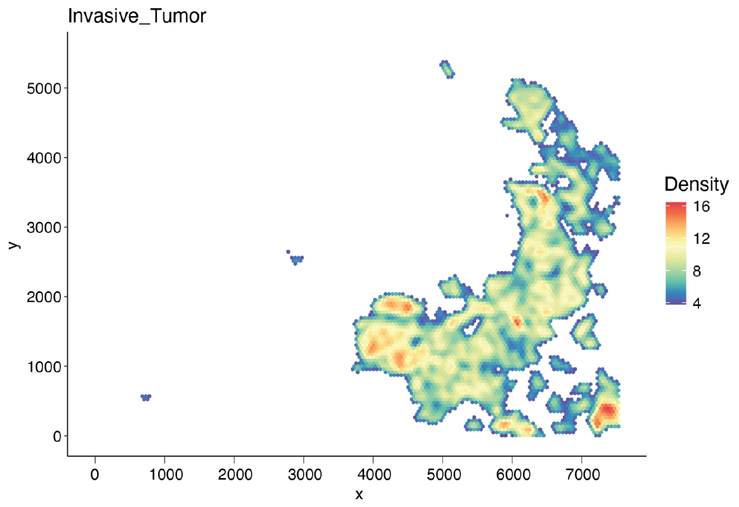


**Figure S6:** Heatmap of spatial density of invasive tumor cells in the Xenium breast carcinoma sample (replicate 1).

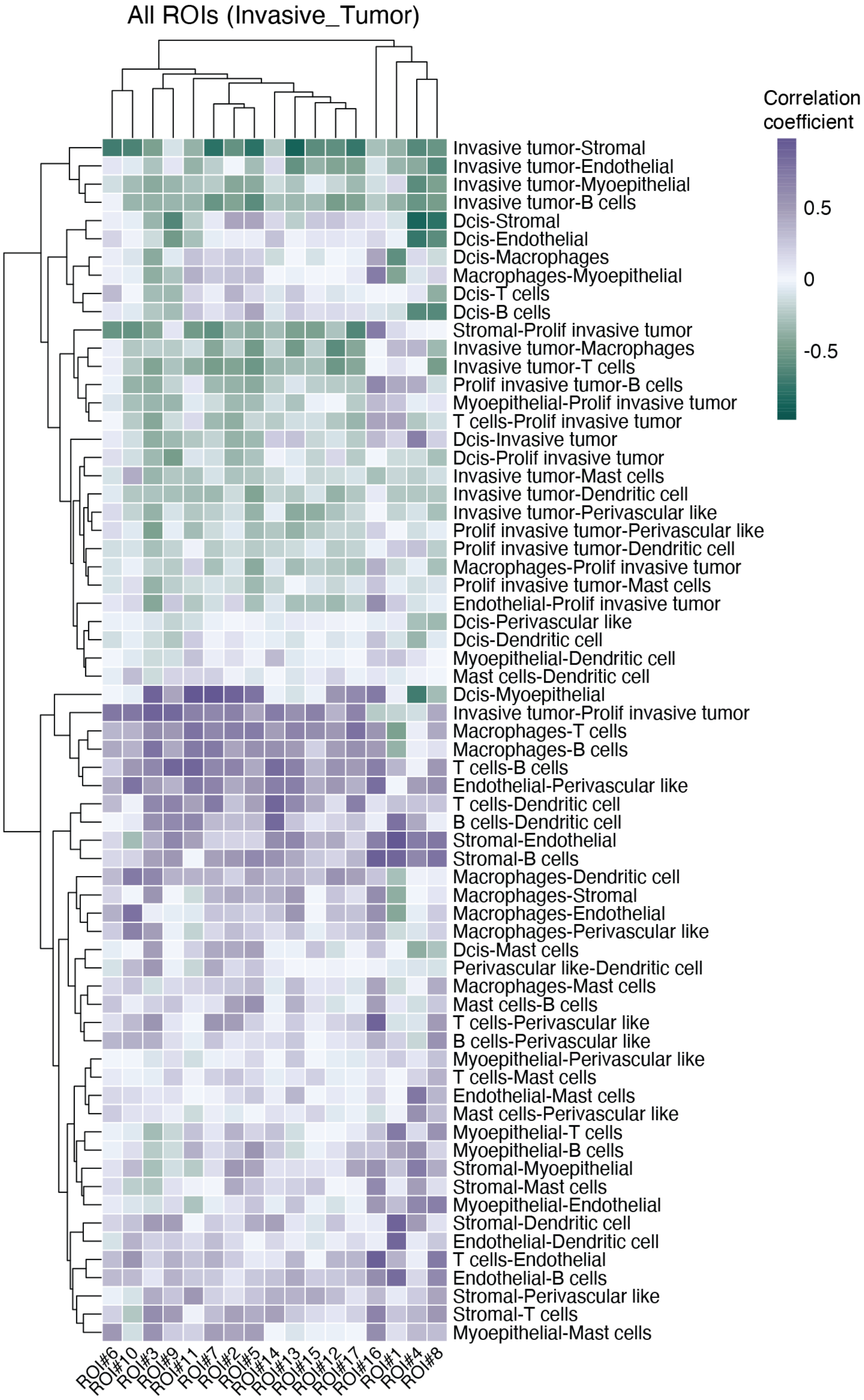


**Figure S7:** Cell type colocalization in all invasive tumor ROIs in the Xenium breast carcinoma sample (replicate 1).

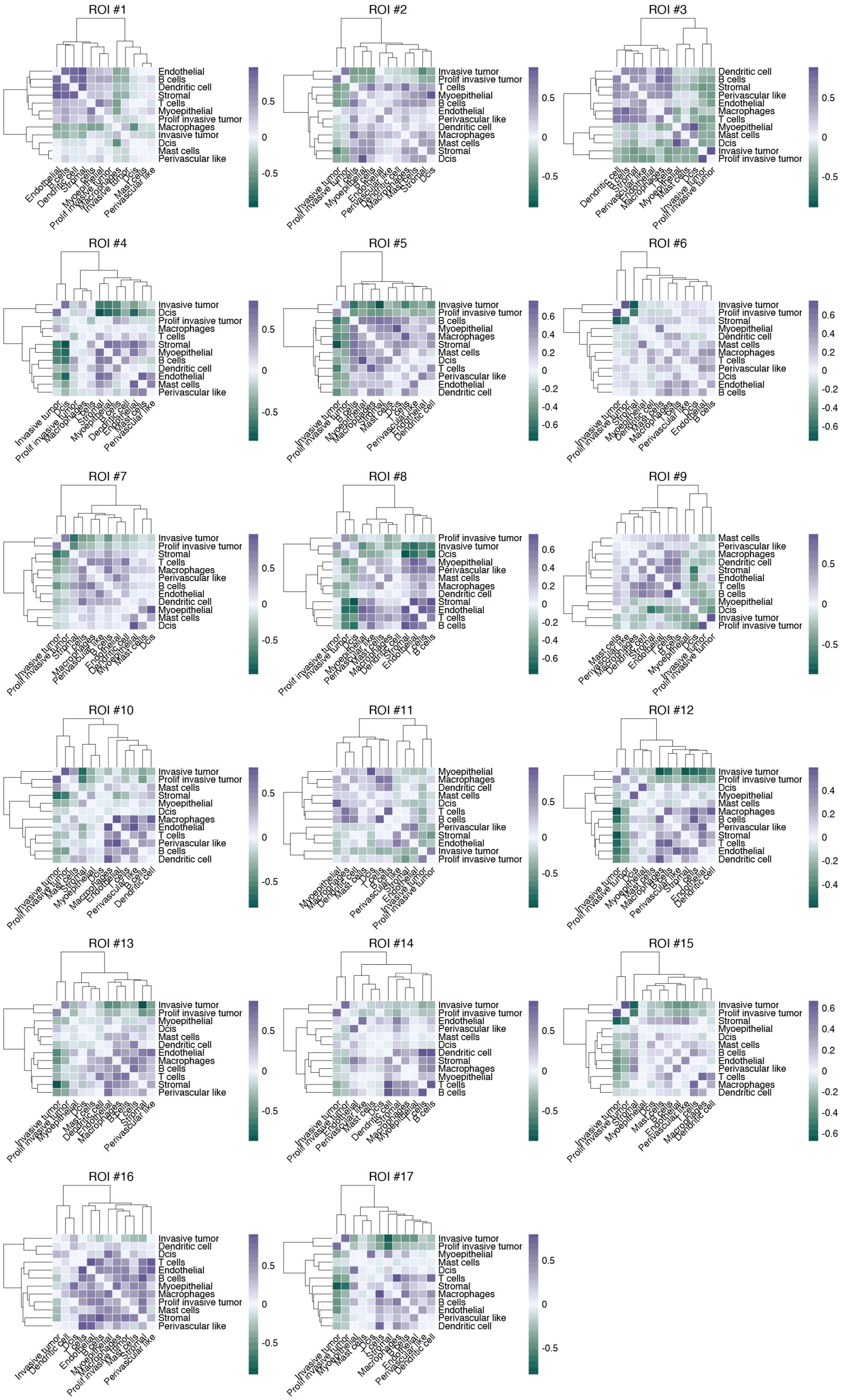


**Figure S8:** Cell type colocalization in each invasive tumor ROI in the Xenium breast carcinoma sample (replicate 1).

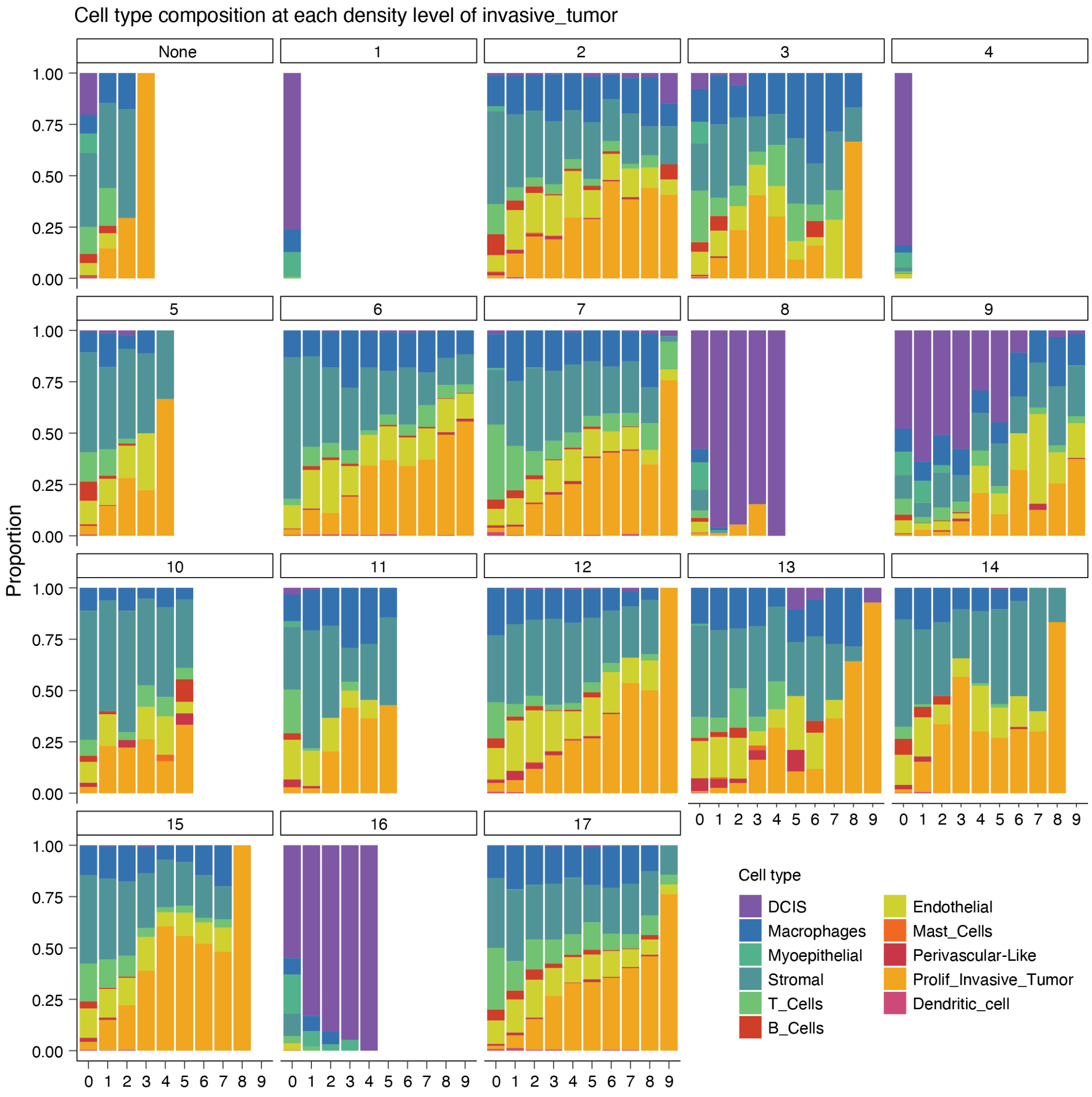


**Figure S9:** Cell type composition at each contour level of invasive tumor density in each invasive tumor ROI in the Xenium breast carcinoma sample (replicate 1).

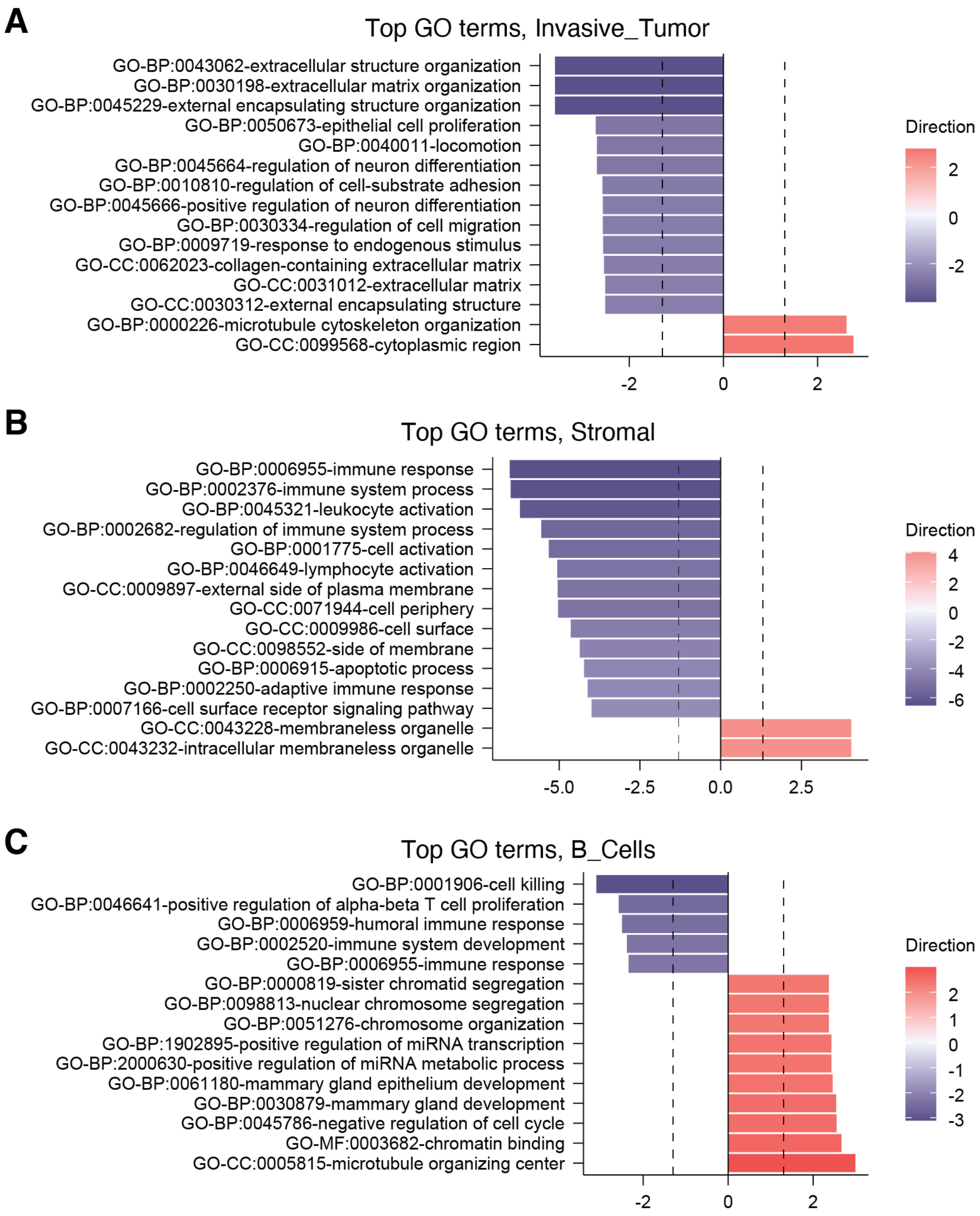


**Figure S10:** Top GO terms associated with invasive tumor densities in invasive tumor cells (A), stromal cells (B) and B cells (C).

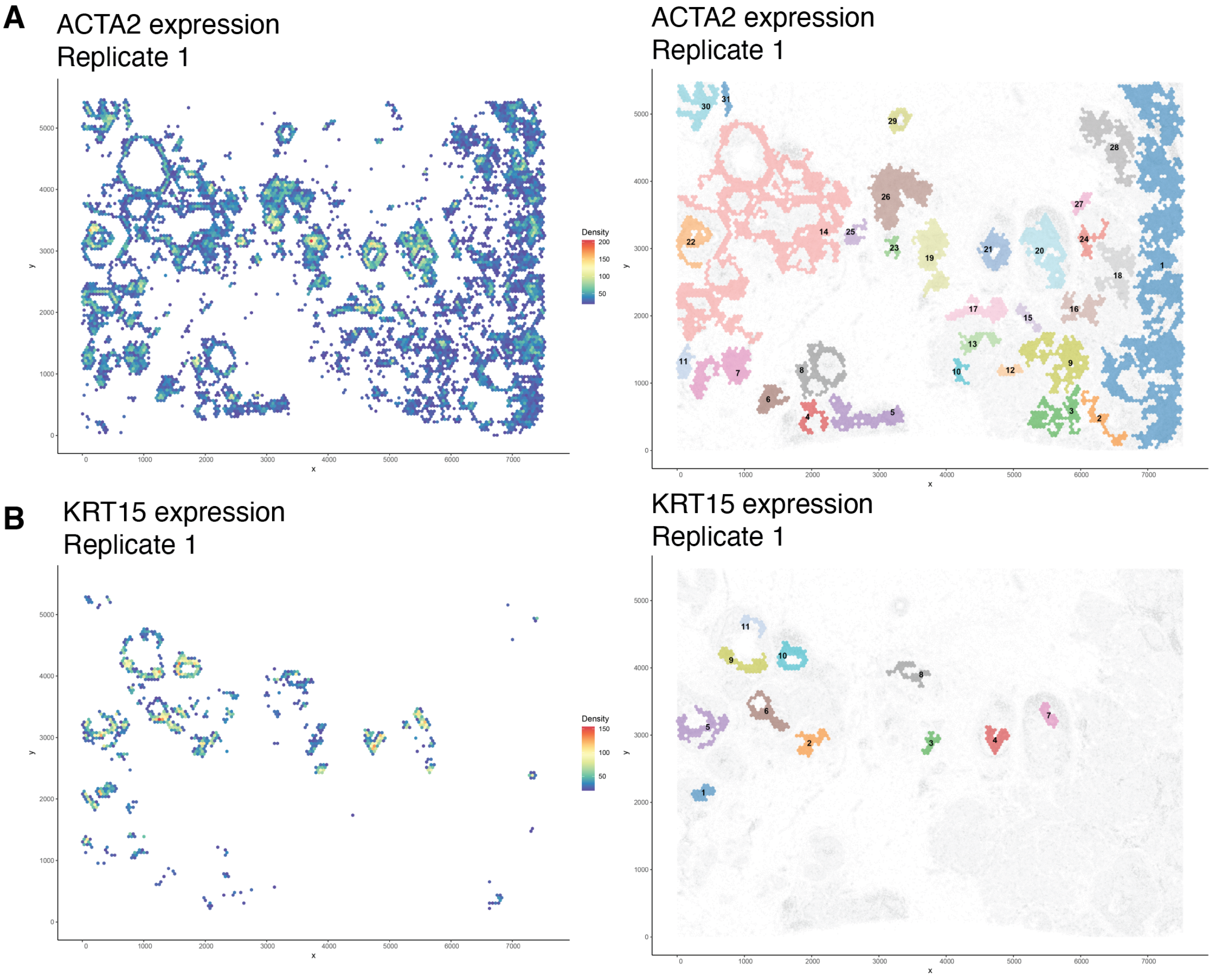


**Figure S11:** Gene-based ROI detection on the Xenium breast carcinoma sample (replicate 1). (A) Spatial density of ACTA2 expression (left) and ROIs detected based on ACTA2 expression overlaied on all cells (right). (B) Spatial density of KRT15 expression (left) and ROIs detected based on KRT15 expression overlaied on all cells (right).

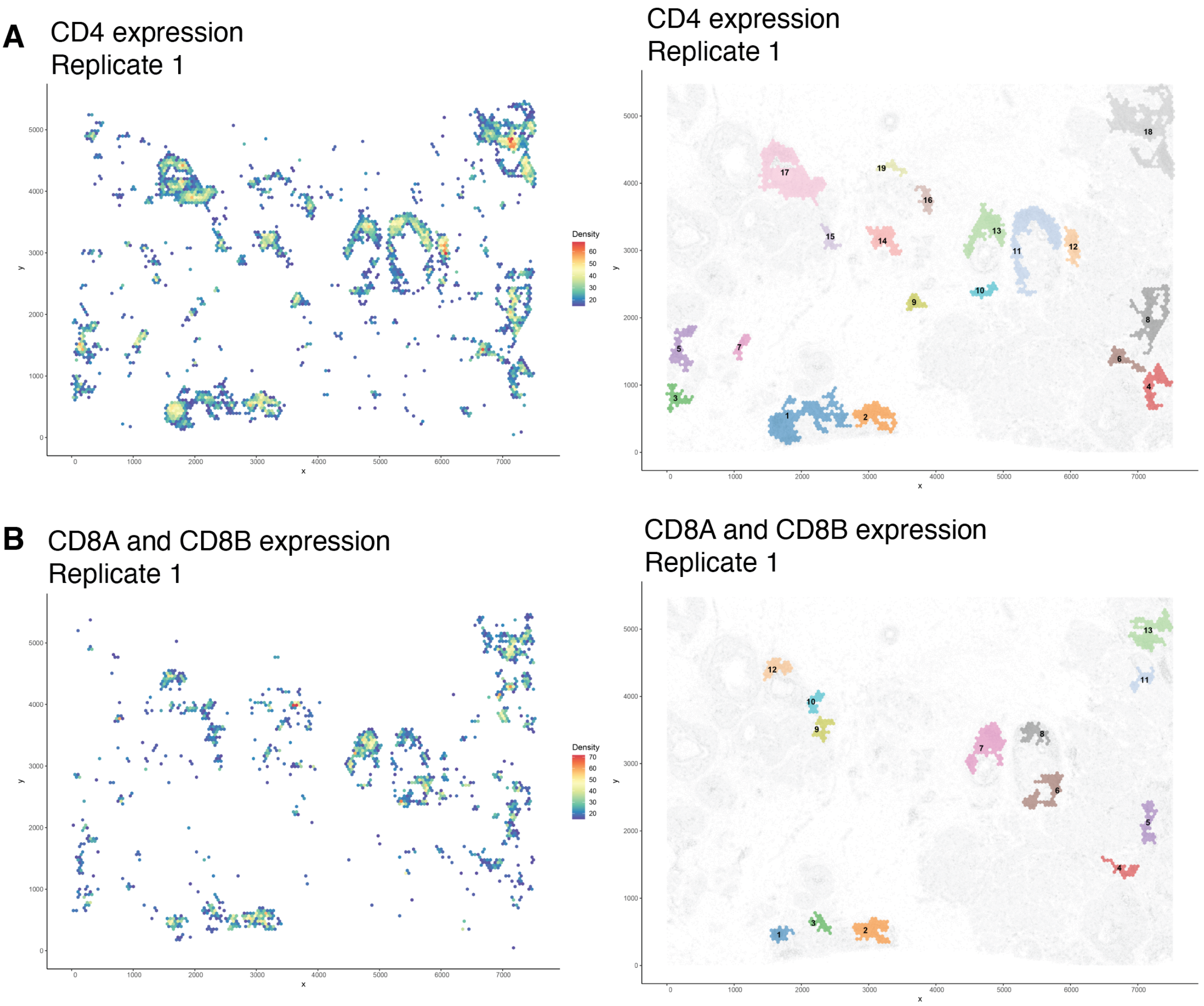


**Figure S12:** Gene- and gene set-based ROI detection on the Xenium breast carcinoma sample (replicate 1). (A) Spatial density of CD4 expression (left) and ROIs detected based on CD4 expression overlaid on all cells (right). (B) Spatial density of CD8A and CD8B expression (left) and ROIs detected based on CD8A and CD8B expression overlaied on all cells (right).
